# Machine learning prediction of cognition from functional connectivity: Are feature weights reliable?

**DOI:** 10.1101/2021.05.27.446059

**Authors:** Ye Tian, Andrew Zalesky

**Author notes:** Corresponding Authors:* Dr Ye Tian, Melbourne Neuropsychiatry Centre, The University of Melbourne, Level 3, Alan Gilbert Building, Victoria, 3053, Australia, *Email:*, Dr Andrew Zalesky, Melbourne Neuropsychiatry Centre, The University of Melbourne, Level 3, Alan Gilbert Building, Victoria, 3053, Australia, *Email:.

## Abstract

Cognitive performance can be predicted from an individual’s functional brain connectivity with modest accuracy using machine learning approaches. As yet, however, predictive models have arguably yielded limited insight into the neurobiological processes supporting cognition. To do so, feature selection and feature weight estimation need to be reliable to ensure that important connections and circuits with high predictive utility can be reliably identified. We comprehensively investigate feature weight test-retest reliability for various predictive models of cognitive performance built from resting-state functional connectivity networks in healthy young adults (n=400). Despite achieving modest prediction accuracies (r=0.2-0.4), we find that feature weight reliability is generally poor for all predictive models (ICC<0.3), and significantly poorer than predictive models for overt biological attributes such as sex (ICC ≈ 0.5). Larger sample sizes (n=800), the Haufe transformation, non-sparse feature selection/regularization and smaller feature spaces marginally improve reliability (ICC<0.4). We elucidate a tradeoff between feature weight reliability and prediction accuracy and find that univariate statistics are marginally more reliable than feature weights from predictive models. Finally, we show that measuring agreement in feature weights between cross-validation folds provides inflated estimates of feature weight reliability. We thus recommend for reliability to be estimated out-of-sample, if possible. We argue that rebalancing focus from prediction accuracy to model reliability may facilitate mechanistic understanding of cognition with machine learning approaches.

## Introduction

Predicting an individual’s cognitive abilities and behavioral traits remains a major goal in neuroscience (Finn et al., 2015; Poldrack et al., 2017; Poldrack et al., 2020; Sui et al., 2020). Facets of human cognition, including intelligence, attention and working memory can be predicted with modest accuracy using machine and deep learning techniques applied to functional magnetic resonance imaging (fMRI) data. Current studies report cross-validated correlations between predicted and actual measures of cognition varying between r=0.1∼0.5, depending on the specific cognitive measure, predictive model and other factors (Chen et al., 2020; Cui and Gong, 2018; Dhamala et al., 2021; Finn and Bandettini, 2021; Finn et al., 2015; Greene et al., 2018; Kong et al., 2019; Li et al., 2019; Mansour et al., 2021; Seguin et al., 2020; Shen et al., 2017). Research efforts are currently focused on improving prediction accuracies through enhanced fMRI modeling, feature engineering, deep learning and larger samples (Abrol et al., 2021; He et al., 2020; Pervaiz et al., 2020; Schulz et al., 2020). As such, prediction accuracy has emerged as one of the most decisive factors differentiating good and bad predictive models of cognitive ability in neuroimaging: My model is better than yours because it is more accurate!

While several predictive models can afford practical utility regardless of whether they are explainable (e.g., prediction of disease outcomes in a clinical setting), accurately predicting intelligence is arguably not an end goal per se. Even if future advances lead to outstanding prediction accuracies, practical, ethical and other considerations may limit use of this technology for often-hyped, real-world applications, such as intelligence testing, cognitive screening tools and similar (Eickhoff and Langner, 2019). A more tangible, realistic and immediate goal of predictive neuroimaging models in cognitive neuroscience is to explain neurobiological processes supporting cognition and to test theoretical cognitive models. While numerous top-down and bottom-up models of cognition have been developed (Kveraga et al., 2007), the brain regions, circuits, networks and dynamic neural processes underlying these models are only partly understood.

A first step toward explaining a machine learning “black box” is through examining feature importance. Feature importance can be quantified using the fitted feature weights (i.e., beta coefficients), usually after appropriate transformation (Haufe et al., 2014). Notwithstanding certain caveats, predictive utility is greatest for features with large weights. As such, neural processes underpinning specific cognitive functions can, in principle, be elucidated and localized to brain regions, connections and networks with large feature weights. However, this requires predictive models that are not only accurate, but also reliable with respect to feature selection and estimation of feature importance. Without reliable feature weights, it is challenging to interpret and explain predictive models, irrespective of how accurate the model can predict a behavior.

Because predictive models are (by definition) validated on independent samples (Rosenberg et al., 2018a), is reproducibility and reliability ensured? Current cross-validation methods are geared toward validating prediction accuracy and can thus furnish confidence intervals quantifying the reliability for accuracy measures, but reliable prediction accuracies do not necessarily imply reliable estimates of feature importance.

Several recent machine learning studies predicting cognitive performance from fMRI-derived brain networks have commented on the test-rest reliability of connectivity feature weights, variably referred to as either consistency, occurrence rate, overlap or consensus (Cui and Gong, 2018; Dhamala et al., 2021; Finn and Bandettini, 2021; Finn et al., 2015; Jiang et al., 2019; Rosenberg et al., 2016). These studies suggest that feature importance estimation is moderately reliable, although reliability has not typically been a core focus of these studies and strong agreement in beta coefficients between cross-validation folds has not always been interpreted in terms of reliable feature weights. Features that overlap between cross-validation folds have been computed to enable succinct visualization of important features and to facilitate external model validation using independent datasets (Rosenberg et al., 2016; Rosenberg et al., 2018b). External validation based on an independent dataset is one of the strongest forms of validation and can provide insight into model generalizability. However, many studies do not have the luxury of an independent/external dataset, and thus model validation has often been performed “internally” on the same dataset using cross-validation schemes. Given that training splits generally do not comprise independent samples between cross-validation folds, this can lead to inflated test-retest reliability estimates, as we demonstrate here. Further work is therefore needed to establish the extent to which the estimation of feature importance and model selection is reliable and reproducible.

If the primary goal is to explain mechanisms underpinning cognition, significance testing may seem a more appropriate instrument than machine learning approaches. However, predictive modelling can alleviate several limitations of conventional statistical inference, including overfitting and *p*-hacking (Rosenberg et al., 2018a; Yarkoni and Westfall, 2017). Predictive and explanatory models are thus complementary, and they can both ultimately assist with elucidating a mechanistic understanding of cognition. Moreover, predictive models can potentially reveal unique insights, given that connectivity features that are important to predicting a cognitive measure do not necessarily correspond with connections that significantly covary with that measure (Bzdok et al., 2020).

The reliability of functional connectivity measurements may impact feature weight reliability. If connectivity features themselves cannot be reliably measured, it is unlikely that feature importance can be reliably estimated. The reliability of resting-state functional connectivity is poor to modest (Noble et al., 2019) and depends on the precise functional connectivity measure, fMRI acquisition length, parcellation scale and many other factors (Birn et al., 2013; Noble et al., 2017; Pannunzi et al., 2017; Shirer et al., 2015; Taxali et al., 2021). The extent to which the reliability (or lack thereof) of functional connectivity measurements impacts the reliability of feature weight estimation remains unclear.

The goal of this study is to evaluate the test-retest reliability of resting-state functional connectivity feature weights estimated by predictive models of intelligence and cognitive function. We consider feature spaces of varying dimensionality, from ∼ 100 connections between broad canonical brain networks, to high-resolution atlas-defined networks comprising more than 10,000 connections. Prediction accuracies and feature weight test-retest reliability are evaluated under several realistic conditions, including different predictive models, varying sample sizes and use of the Haufe transformation. We also compare the reliability of univariate statistics and investigate whether predictive models of overt biological attributes (i.e., sex) yield more reliable feature weights than cognitive models. Finally, we provide recommendations for maximizing feature weight reliability and elucidate a tradeoff between reliability and prediction accuracy. We emphasize that our conclusions and recommendations do not necessarily generalize to predictive models of diagnostic status, disease outcomes and other clinical variables. Here, we focus on predictive modeling of cognition in healthy young adults. We hope that our work encourages researchers to consider both feature weight reliability and prediction accuracy when evaluating predictive models of cognitive performance.

## Material and methods

### Participants and neuroimaging data

Minimally preprocessed resting-state functional magnetic resonance imaging (fMRI) data were sourced from the Human Connectome Project (HCP) S1200 release (Van Essen et al., 2013). Participants were young healthy adults (n=1113) aged between 22 and 37 years. Two sessions of resting-state fMRI (REST1 and REST2) were acquired for each participant on two consecutive days, where each session comprised two runs (right-to-left and left-to-right phase encoding) of 14m33s each (TR=720ms, TE=33.1ms, voxel dimension: 2×2×2 mm^3^). All images were acquired on a customized Siemens Skyra 3 Tesla MR scanner using a multiband echo planar imaging sequence. Further image acquisition details can be found elsewhere (Smith et al., 2013). Participants who completed all four runs and cognitive assessments were included in this study, yielding a final sample of 958 individuals (mean age 28.7±3.7 years, 453 males).

The minimal preprocessing pipeline included removal of spatial artifacts and distortions, correction of head motion and spatial registration to the MNI (Montreal Neurological Institute) standard space, as described in detail elsewhere (Glasser et al., 2013). Data were analyzed in CIFTI (Connectivity Informatics Technology Initiative) format. Cortical data were projected onto a standard surface mesh (fs_LR) comprising ∼32k vertices in each hemisphere, using a multimodal surface matching approach, referred to as MSM-ALL (Robinson et al., 2014). Subcortical data remained in volumetric format and were spatially aligned to the MNI space using the FNIRT nonlinear registration algorithm (Woolrich et al., 2009). The data were then spatially smoothed with surface and parcel constrained smoothing of 2mm FWHM (full width at half maximum). Motion-related artifacts and structured physiological noise were removed with ICA-FIX (Griffanti et al., 2014; Salimi-Khorshidi et al., 2014). In addition, Wishart filtering, a PCA-based data denoising method, was performed on the denoised fMRI time series (Glasser et al., 2018; Glasser et al., 2016a; Glasser et al., 2016b) to further improve the signal-to-noise ratio. Head motion was quantified using framewise displacement (Power et al., 2012) and was included as a covariate in all statistical analyses.

### Functional connectivity estimation

The preprocessed fMRI time series were temporally concatenated across the four runs, yielding approximately one hour of data for each individual. Concatenation of the four runs ensured sufficient data to enable stable estimates of functional connectivity (Birn et al., 2013; Gordon et al., 2017; Gratton et al., 2018). Whole-brain functional connectivity matrices were mapped using established cortex (Glasser et al., 2016a) and subcortex atlases (Tian et al., 2020). Specifically, fMRI signals were averaged across all vertices and voxels comprising each cortical (N=360) and subcortical (N=16) region, respectively. The Pearson correlation coefficient was used to estimate the temporal dependence between each pair of regional time series, yielding a symmetric functional connectivity matrix of dimension 376 × 376 for each individual. The connectivity matrix was r-to-z transformed (Fisher transformation), followed by vectorization of the upper triangle, resulting in (376 × 375)/2=70,500 connectivity features. While the Pearson correlation coefficient is the most widely used measure of functional connectivity, alternative measures may yield improved prediction models (Pervaiz et al., 2020).

In complementary analyses, connectivity features were also derived from whole-brain functional networks previously mapped using independent component analysis (ICA) and a dual regression approach (Beckmann and Smith, 2004; Filippini et al., 2009). Spatial scales comprising 15, 24, 50, 100, 200 and 300 components (nodes) were mapped, yielding feature spaces ranging between 105 and 31,350 functional connections for the current study. Further details about the ICA-based networks are available as part of the HCP S1200 PTN data release: https://www.humanconnectome.org/storage/app/media/documentation/s1200/HCP1200-DenseConnectome+PTN+Appendix-July2017.pdf

### Cognitive measures

Cognitive performance was measured using: i) *fluid intelligence* (fIQ); ii) *crystalized intelligence* (cIQ); and iii) an overall composite measure of cognition (IC-Cognition). Measures of fluid and crystallized intelligence acquired by the HCP were used without modification. Fluid intelligence is one of the most extensively studied cognitive phenotypes in the neuroimaging literature (Finn et al., 2015). It was measured in the HCP using the Penn Matrix Test, an abbreviated version of the Raven’s Progressive Matrices test, with 24 items (Bilker et al., 2012). Crystalized intelligence was measured using the NIH Toolbox Picture Vocabulary Test, which assesses general vocabulary, providing an indicator of crystalized ability (Weintraub et al., 2013). A composite measure of overall cognition was derived using independent component analysis applied to 109 behavioral items, as described elsewhere (Tian et al., 2020). This provided a single continuous summary score of overall cognitive performance across a range of tasks and behaviors, referred to as *IC-Cognition* in the current study.

### Cross-validation

Cross-validated predictive models were trained to predict fIQ, cIQ and IC-Cognition. A half-split cross-validation procedure was designed to estimate the test-retest reliability of the feature weights (i.e., beta coefficients) constituting each of these models (Figure 1). The cross-validation procedure controlled for genetic relatedness, as defined by the 420 families among the 958 individuals in the final sample. Specifically, to ensure independence of the test and train sets, we randomly selected one individual from each family among a random set of 400 families, resulting in a group of 400 genetically unrelated individuals. The selected 400 individuals were further subdivided into two subgroups of 200 each, defining the train and/or test set, respectively. To minimize sampling biases, this sampling and half-split procedure was repeated 100 times, resulting in 100 pairs of independent train-test data splits. Leaving out 20 families from each random sample ensured that individuals who were the sole member of a family (n=90) were not selected in all 100 train-test pairs. Sex ratios did not significantly differ between half-split pairs (*χ*^2^=0-5.7, p>0.05, FDR corrected across 100 pairs).

**Figure 1.**
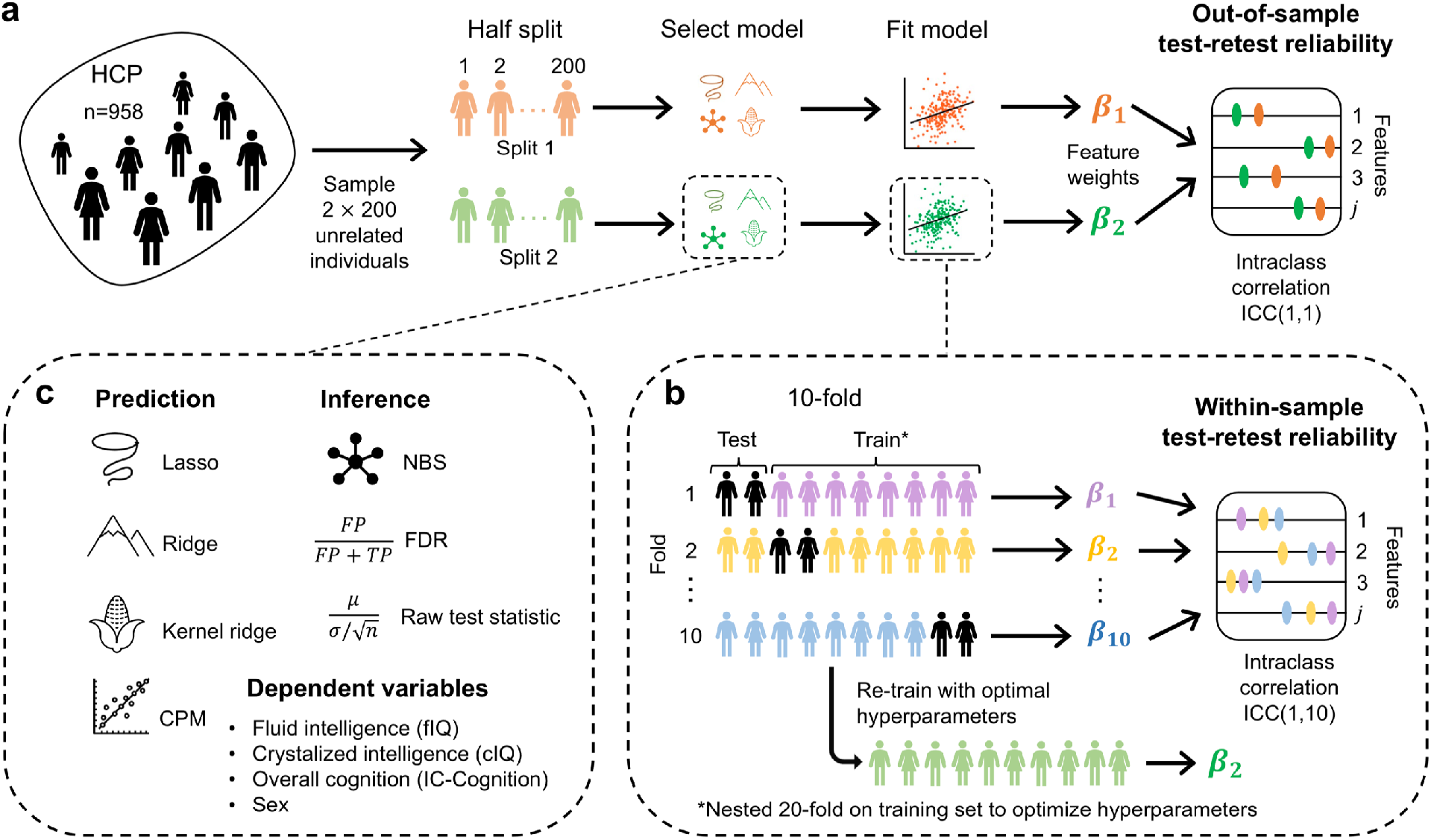
Cross-validation procedure to estimate feature weight test-retest reliability and catalogue of predictive models. **(a)**, For each half-split cross-validation iteration, 400 genetically unrelated individuals were randomly selected from 958 individuals. The 400 individuals were split into two folds (Split 1 and Split 2) to define the train and/or test set, respectively. Models were independently trained on each of two half splits to predict fIQ, cIQ and IC-Cognition, yielding two sets of beta coefficients (***β***_2_ and ***β***_1_) for each cognitive measure. The intraclass correlation coefficient (ICC) between the two sets of beta coefficients provided an out-of-sample estimate of feature weight test-retest reliability. **(b)**, Test-retest reliability is more commonly estimated based on agreement in feature weights across cross-validation folds. Within each data split (Split 1 or Split 2), 10-fold cross-validation yielded 10 sets of beta coefficients (***β***_2_ … ***β***_2*n*_) estimated from each of 10 training sets. The ICC across the 10 sets of beta coefficients provided a within-sample estimate of feature weight test-retest reliability. **(c)**, Predictive models investigated included least absolute shrinkage and selection operator (lasso), ridge, kernel ridge regression and connectome-based predictive modeling (CPM). As a comparison, test-retest reliability was also estimated for inferential statistical methods, including univariate statistics for each connection as well as significant connections identified by the network-based statistic (NBS) and the false discovery rate (FDR).

#### Out-of-sample test-retest reliability

For each train-test data split, predictive models were trained using one of the two splits to predict fIQ, cIQ and IC-Cognition. The trained model was then applied to the other split (test set) to evaluate prediction accuracy. The same procedure was repeated twice to ensure that each split was treated as a train and test set separately. This yielded two sets of beta coefficients (***β***_1_ and ***β***_2_), one for each of the two data splits (Split 1 and Split 2), consistent with two-fold cross-validation. The intraclass correlation coefficient (ICC), a commonly used measure for test-retest reliability estimation in fMRI (Noble et al., 2019; Noble et al., 2021) was used to quantify the extent of consistency in beta coefficients between the two splits. The ICC formulation originally proposed by Fisher was used, as given by,

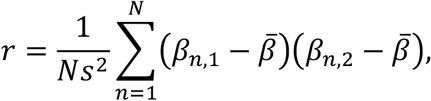

where *N* is the number of features, *β*_*n*,1_ is the feature weight for the nth feature estimated in the first split and analogously for *β*_*n*,2_. In this formulation, *s*^2^ and 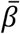 are pooled estimates of the variance and mean, respectively, defined as,

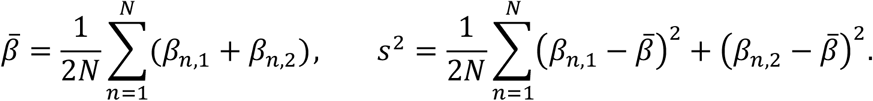

This is sometimes referred to as the 1-1 formulation of the ICC. The ICC is sometimes conceptualized in terms of the extent of agreement in measurements made by a set of observers measuring the same set of objects. In the above formulation, each connection (i.e., feature weight) corresponds to a separate “object”, while the two “observers” are Split 1 and Split 2. This differs from many other neuroimaging studies using the ICC, where “objects” typically correspond to different subjects and “observers” are subject measurements taken across different days/sessions (Noble et al., 2019). Given that our aim is to assess the consistency in feature weights between half split pairs, this is the most appropriate formulation here because our “objects” of interest (i.e., feature weights) are not measured at the level of individual subjects.

Before computing *r*, the feature weight vectors ***β***_1_ and ***β***_2_ were each z-scored, rendering the ICC equivalent to the Pearson correlation coefficient (i.e., 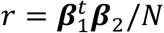 because 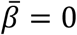 and *s*^2^ = 1 after z-scoring). Without z-scoring, ICC values were marginally lower in rare cases, but otherwise unchanged.

Given that individuals comprising the two data splits were completely independent and the model was independently trained in each data split, the estimated consistency in beta coefficients between the two data splits is referred to as *out-of-sample test-retest reliability* (Figure 1a). High ICC values indicate that feature importance can be reliably localized to specific functional connections.

#### Within-sample test-retest reliability

In contrast to out-of-sample estimation, the consistency in feature selection (i.e., test-retest reliability of feature weights) is more commonly evaluated by i) averaging (or summing) feature weights across k-fold or leave-one-out cross-validation iterations (Chen et al., 2020; Dhamala et al., 2021; Greene et al., 2018); and/or, ii) selecting features with beta coefficients exceeding a certain threshold in a proportion of all cross-validation training folds (Finn and Bandettini, 2021; Finn et al., 2015; Jiang et al., 2019; Rosenberg et al., 2016; Shen et al., 2017). The first (averaging across training sets) and second approaches (consensus across training sets) both provide within-sample measures of reliability because the training sets are not mutually exclusive between folds, except for the case of two-fold cross-validation. The lack of mutual exclusivity between trainings sets is greatest for leave-one-out cross-validation, potentially resulting in inflated estimates of test-retest reliability.

As exemplified in Figure 1b, k-fold (k=10) cross-validation was performed in each of the two data splits (Split 1 and Split 2). Each fold was treated as a test set once and the remaining 9 folds were used to train the prediction model. This yielded 10 sets of beta coefficients (***β***_1_ … ***β***_10_). *Within-sample test-retest reliability* was defined as the ICC across the 10 sets of beta coefficients. For *out-of-sample test-retest reliability*, the 10 sets of beta coefficients were first averaged within each data split, consistent with previous studies (Chen et al., 2020; Dhamala et al., 2021; Greene et al., 2018), and the ICC was computed between the two averaged beta coefficients. We also computed within-sample and out-of-sample reliability using leave-one-out cross-validation, enabling comparison with previous studies (Finn et al., 2015; Jiang et al., 2019; Rosenberg et al., 2016; Shen et al., 2017).

### Predictive models and prediction accuracy

Four commonly used linear regression models were trained to predict fIQ, cIQ and IC-Cognition (Figure 1c): i) the least absolute shrinkage and selection operator (lasso) (Tibshirani, 1996); ii) ridge regression (Hoerl and Kennard, 1970); iii) kernel ridge regression (Li et al., 2019); and, iv) connectome-based predictive modelling (CPM) (Shen et al., 2017). We did not consider deep neural networks and nonlinear models because they do not allow for straightforward decoding of the relationship between predictive features and the target variable of interest. Regularization and model training are described in Supplementary Materials. In brief, a nested 20-fold cross-validation was used for hyperparameter optimization to minimize the cross-validation error in each training split for lasso, ridge and kernel ridge regression. Least-squares regression was used for continuous cognitive variables, whereas logistic regression was used for sex prediction. The hyperparameters resulting in the smallest error averaged across all inner test folds were selected to compute the beta coefficients, using all individuals in the outer training split. Similarly, nested 20-fold cross-validation was used for CPM to find the optimal *p*-value threshold to maximize the positive or negative association between the summation of functional connectivity strengths and the cognitive measure.

Individual variation in age, sex and head motion (FD) was regressed from each of the three cognitive measures before model training. Beta coefficients for these confounds were estimated in the training set and then applied to the test set to avoid leakage. No confound regression was performed prior to sex prediction to preserve the binary nature of this variable.

Following previous studies (Finn et al., 2015; Li et al., 2019), prediction accuracy was quantified based on the Pearson correlation coefficient between the observed and the out-of-sample predicted cognitive scores across all individuals in the test set. For sex, FD as well as age were controlled when evaluating prediction accuracy (partial correlation), given the significant age difference between males (mean age 27.9±3.7 years) and females (mean age 29.5±3.6 years) in the sample (t=6.80, p=1.82×10^−11^).

Prediction accuracy and test-retest reliability was computed separately in each pair of the 100 train-test data splits, as described above, resulting in 100 out-of-sample and 200 within-sample (100 per half split) estimates of feature weights test-retest reliability and 200 out-of-sample prediction accuracy estimates for each cognitive variable and sex. For kernel ridge regression, the feature space comprised interindividual similarity in connectivity matrices between individuals in the training set. Connectivity feature weights were thus computed as the weighted mean of functional connectivity values across individuals in the training set, where individuals were weighted by their beta coefficients (Chen et al., 2020).

### Haufe transformation

The Haufe transformation (Haufe et al., 2014) was applied to the beta coefficients before evaluating test-retest reliability. This transformation improves the interpretability of feature weights and ensures that important features are weighted highly. The Haufe-transformed beta coefficient for a given connection was computed as *β*^*Haufe*^ = *ρ*^*T*^ *ŷ*/*N*, where *ρ* is the *N* × 1 vector of standardized functional connectivity values and *ŷ* is the *N* × 1 vector of predicted cognitive scores for each of the *N* individuals in the training set. The Haufe transformation was not used for CPM (Shen et al., 2017).

### Randomization

Cognitive measures were randomly permuted across individuals, thereby randomizing associations between cognition and functional connectivity. Randomization was performed independently for each data split and the predictive models were retrained using the randomized data. The test data within each split was not randomized. This yielded 100 samples of test-retest reliability and 200 samples of prediction accuracy to establish chance-level expectations. Two-sample t-tests were used to assess whether the observed prediction accuracies and reliability were significantly greater than chance-level expectations. The false discovery rate (FDR) was controlled at a threshold of 5% across all predictive models for each cognitive variable and sex.

## Results

We sought to determine whether predictive utility (i.e., feature importance) can be reliably assigned to resting-state functional connectivity features comprising predictive models of cognitive performance. To this end, we evaluated the test-retest reliability of feature weights estimated using predictive models of fluid intelligence (fIQ), crystallized intelligence (cIQ), and overall cognitive performance (IC-Cognition) in a cohort of healthy young adults. Samples of 400 (or 800) individuals were repeatedly drawn from the cohort and split into halves (i.e., half-split cross-validation; see Methods). Various machine learning approaches were used to train predictive models for each half-split and the intraclass correlation coefficient (ICC) was used to evaluate feature weight test-retest reliability between the two halves.

We found that fIQ, cIQ and IC-Cogntion can be predicted with modest accuracy (r=0.2-0.4) using resting-state functional connectivity strengths (Figure 2a), consistent with previous literature (Chen et al., 2020; Dhamala et al., 2021; Finn et al., 2015; Li et al., 2019; Mansour et al., 2021; Seguin et al., 2020). However, estimated feature weights consistently showed poor reliability (ICC<0.3) across all machine learning models and measures of cognitive performance (Figure 2b). In comparison, feature weights for sex prediction showed moderate reliability (ICC>0.6) for several models (Figure 2b). Therefore, unlike an overt biological attribute such as sex, cognitive performance could not be reliably localized to specific connections defining the most important features of a predictive model. This motivated investigation of whether feature weight test-retest reliability could be improved by using mass univariate approaches, larger sample sizes, larger/smaller feature spaces and/or more sophisticated predictive models. But first, we investigated prediction accuracy and feature weight reliability in more detail for the nominal sample size of 400 individuals (200 per half-split) and a feature space comprising 70,500 resting-state functional connections.

**Figure 2.**
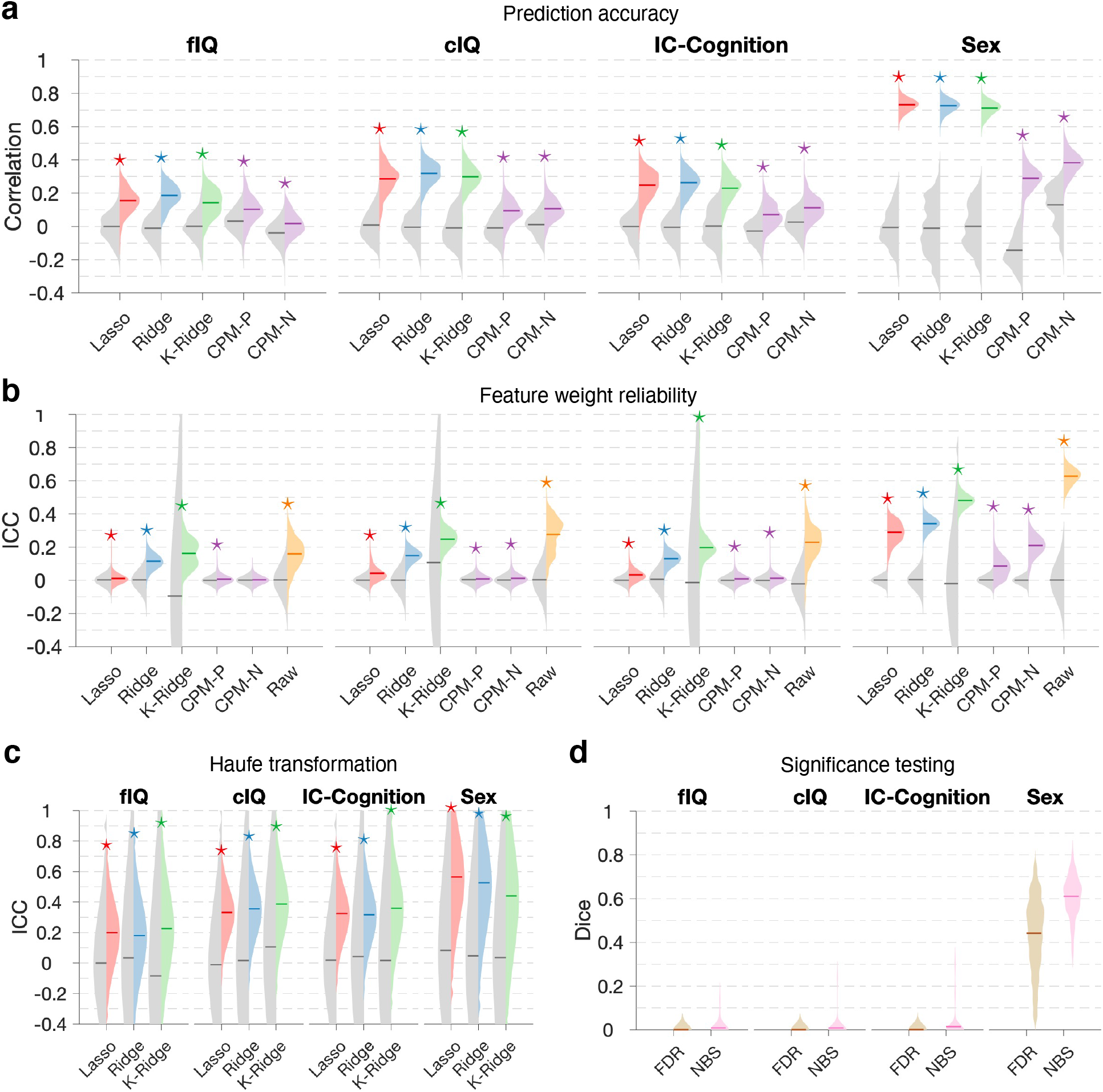
Prediction accuracy and feature weight test-retest reliability estimated using half-split cross-validation in 400 unrelated individuals. **(a)**, Pearson correlation coefficient between observed and out-of-sample predicted fluid intelligence (fIQ), crystalized intelligence (cIQ), overall cognition (IC-Cognition) as well as sex (partial linear correlation, 0-female; 1-male). Violin plots show the distribution of correlation coefficients across 100 half-split pairs (left: chance; right: observed). Asterisks are shown to indicate mean Pearson correlation coefficients significantly exceeding chance-level predictions (two-sample t-test, p<0.05, false discovery rate (FDR) correction across five models). Violin plots are shown for each predictive model; namely, lasso regression, ridge regression, kernel ridge regression and connectome-based predictive modeling (CPM). CPM-P, positive associations; CPM-N, negative associations. **(b)**, Feature weight test-retest reliability (out-of-sample) quantified with the intraclass correlation coefficient (ICC). Raw refers to the univariate test statistic for each connection, which assesses the null hypothesis of an absence of association between cognitive performance and functional connectivity strength. Violin plots show the distribution of ICC values across 100 half-split pairs (left: chance; right: observed). Asterisks are shown to indicate ICC mean values significantly greater than chance (two-sample t-test, p<0.05, FDR correction across five models). **(c)**, Same as (b), but with the Haufe transformation applied to feature weights before evaluating test-retest reliability. **(d)**, The network-based statistic (NBS) and false discovery rate (FDR) were used to identify connections with functional connectivity strengths that significantly correlated with cognitive performance (family-wise error rate and FDR controlled at 0.05, respectively). Violin plots show the distribution of dice coefficients quantifying overlap in connections declared statistically significant across 100 half-split pairs. Central mark on each violin plot indicates the mean value.

### Prediction accuracy

As shown in Figure 2a, correlation coefficients between predicted and actual cognitive performance significantly exceeded chance-level predictions for all cognitive measures: fIQ (mean ± standard deviation (SD), lasso: r=0.16 ± 0.07; ridge: r=0.19 ± 0.07; kernel ridge: r=0.14 ± 0.09; CPM-positive: r=0.10 ± 0.08; CPM-negative: r=0.02 ± 0.06), cIQ (Lasso: r=0.29±0.09; ridge: r=0.32±0.07; kernel ridge: r=0.30±0.07; CPM-positive: r=0.09±0.07; CPM-negative: r=0.11±0.07), IC-Cognition (lasso: r=0.25±0.08; ridge: r=0.26±0.07; kernel ridge: r=0.23±0.08, CPM-positive: r=0.07±0.07; CPM-negative: r=0.11±0.07). Sex could be predicted with greater accuracy than cognitive performance (lasso: r=0.73 ± 0.03; ridge: r=0.73 ± 0.03; kernel ridge: r=0.71 ± 0.04; CPM-positive: r=0.29 ± 0.07; CPM-negative: r=0.38±0.07).

### Feature weight test-retest reliability

Feature weight test-retest reliability was evaluated using ICC. ICC was computed between pairs of feature weights (i.e., beta coefficients) across 100 half-split pairs, yielding 100 ICC values. As shown in Figure 2b, ICC values significantly exceeded chance-level expectations for all cognitive measures: fIQ (mean± SD, lasso: ICC=0.01±0.02; ridge: ICC=0.11±0.04; kernel ridge: ICC=0.16±0.08; CPM-positive: ICC=0.007±0.02), cIQ (lasso: ICC=0.04±0.04; ridge: ICC=0.15±0.04; kernel ridge: ICC=0.25±0.06; CPM-positive: ICC=0.008±0.02; CPM-negative: ICC=0.01±0.02), IC-Cognition (lasso: ICC=0.03±0.03; ridge: ICC=0.13±0.04; kernel ridge: ICC=0.20 ± 0.09; CPM-pos: ICC=0.009 ± 0.02; CPM-negative: ICC=0.01 ± 0.03). While significantly greater than chance, feature weight test-retest reliability was poor for all three cognitive measures (ICC<0.3), regardless of the predictive model, and substantially lower than the feature weight reliability of connectivity features predicting sex (lasso: ICC=0.29±0.06; ridge: ICC=0.34±0.04; kernel ridge: ICC=0.48±0.03; CPM-positive: ICC=0.09±0.07; CPM-negative: ICC=0.21±0.06). Of the predictive models considered, ICC was highest for kernel ridge regression, but use of a kernel also increased the variability of ICC values across half-split pairs.

Poor feature weight reliability was not due to stochasticity in hyperparameter optimization and model fitting. Repeated model fitting with random initial conditions in the same set of individuals yielded highly consistent beta coefficients for lasso and ridge regression (ICC>0.99 across 20 repetitions). Moreover, feature weights for lasso and ride regression are unique and well-defined when all predictors are continuous variables (Tibshirani, 2013), such as functional connectivity strengths.

### Haufe transformation

We next investigated the impact of the Haufe transformation on feature weight test-retest reliability. This transformation is often applied to feature weights to improve their interpretability (Haufe et al., 2014). We found that while transformation improved ICC values between half-split pairs on average, ICC variability markedly increased (Figure 2c). ICC values once again significantly exceeded chance-level expectations for all cognitive measures: fIQ (mean± SD, lasso: ICC=0.20±0.21; ridge: ICC=0.18±0.25; kernel ridge: ICC=0.23±0.30), cIQ (lasso: ICC=0.33±0.16; ridge: ICC=0.36±0.18; kernel ridge: ICC=0.39±0.21), IC-Cognition (lasso: ICC=0.32±0.17; ridge: ICC=0.32±0.19; kernel ridge: ICC=0.36±0.25). Feature weight reliability for sex prediction also benefited from transformation (lasso: ICC=0.56±0.30; ridge: ICC=0.53±0.32; kernel ridge: ICC=0.44±0.39). We conclude that the Haufe transformation can improve feature weight reliability *on average*, but increased variability following transformation can potentially result in unpredictable performance.

### Mass univariate significance testing

Having found relatively poor feature weight test-retest reliability, we next investigated whether mass univariate significance testing would enable more reliable inference than predictive modeling. A test statistic and corresponding uncorrected *p*-value was independently computed for each connection to test the null hypothesis of an absence of association between functional connectivity strength and cognitive performance. The network-based statistic (NBS) (Zalesky et al., 2010) and false discovery rate (FDR) were then used to correct for multiple testing across the set of 70,500 connections, controlling the family-wise error rate and FDR at 0.05, respectively (NBS primary threshold: t-statistic=2; 5,000 permutations). These methods were repeated for 100 half-split pairs, using the same random sampling procedure described above to generate each pair. The Dice coefficient was then used to evaluate the extent of overlap in significant connections between each half-split pair, analogous to the use of ICC for feature weights. This yielded 100 Dice coefficients for each cognitive measure. As shown in Figure 2d, we found that Dice values were exceedingly small for all cognitive measures, whereas sex differences showed moderate Dice values between half-split pairs (FDR: 0.44±0.17; NBS: 0.61±0.09). Furthermore, the proportion of half-split pairs for which the null hypothesis was rejected for at least one connection was relatively small for all cognitive measures: fIQ (FDR: 24/200; NBS: 32/200; 100 half-split pairs = 200 splits), cIQ (FDR: 26/200; NBS: 32/200) and IC-Cognition (FDR: 36/200; NBS: 36/200). In contrast, the null hypothesis was rejected for all half-split pairs when testing for sex differences (FDR: 200/200; NBS: 200/200). We conclude that mass univariate significance testing does not enable more reliable inference about connections governing cognitive performance than predictive modeling.

We also investigated the test-retest reliability of the univariate test statistic (i.e., t-statistic) computed for each connection during mass univariate significance testing. Test-retest reliability was once again evaluated using the ICC between half-split pairs. We found that the univariate statistics showed the greatest test-retest reliability in the current study (Figure 2b, violin plots labeled “Raw”). This suggests that dichotomization based on a statistical significance threshold or categorical feature selection (i.e., lasso) is detrimental to reliability. However, it is important to note that the improvement in test-retest reliability achieved by avoiding dichotomization was modest and the ICC remained below 0.4 for all cognitive measures.

### Impact of sample size

While the sample size (n=400) is comparable or larger than many neuroimaging studies (Finn et al., 2015; Greene et al., 2018; Jiang et al., 2019; Li et al., 2019; Liégeois et al., 2019; Varoquaux, 2018) that have investigated predictive models of cognition, we next aimed to test whether prediction accuracy and test-retest reliability would improve for larger sample sizes. To this end, we doubled the sample size (n=800, with 400 per half-split) and repeated the above experiments (Supplementary Figure 1). While it was no longer possible to ensure that all 800 individuals were genetically unrelated, members of the same family were allocated to either the train or test set, but not both. In Figure 3, the left (n=800) and right (n=400) lobes of each violin plot compare prediction accuracies and feature weight test-retest reliabilities between the two sample sizes. For most predictive models, prediction accuracy (Figure 3a) and feature weight test-retest reliability (Figure 3b) significantly improved for n=800 compared to n=400, although the improvement was modest in most cases, and was most prominent for sex prediction. Doubling the sample size also marginally improved the Haufe-transformed feature weight reliability (Figure 3c) and the reliability of mass univariate significance testing (Figure 3d), although these improvements were only significant for sex prediction. Taken together, these results suggest that substantial increases in sample size lead to relatively modest improvements in the reliability of feature weights.

**Figure 3.**
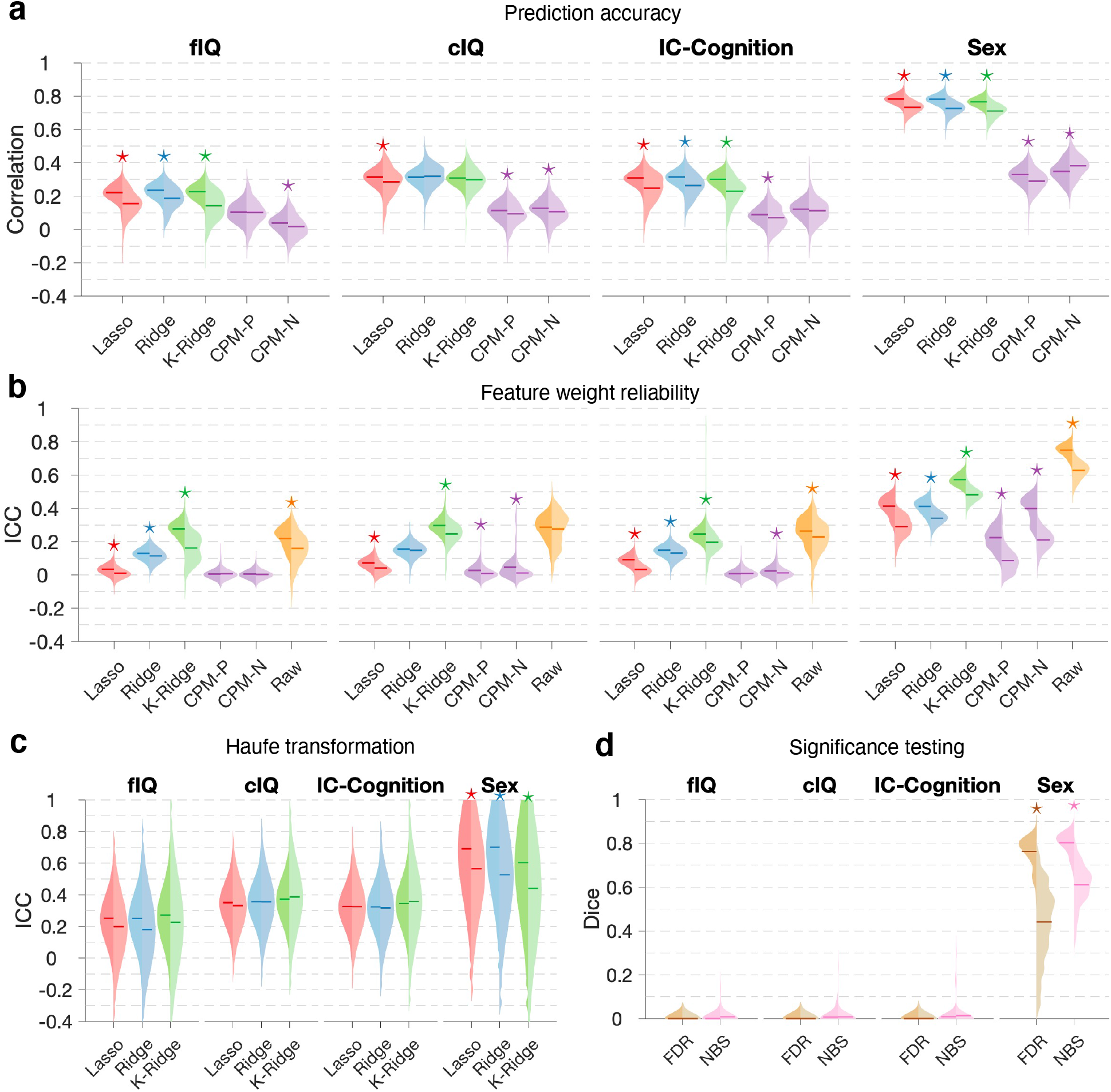
Comparison of half-split cross-validation using 400 and 800 individuals. Figure layout is the same as Figure 2, except the two lobes of each violin plot now compare sample sizes of 800 (left lobe, 400 per half-split) and 400 (right lobe, 200 per half-split). Asterisks are shown to indicate significant differences in mean ICC and Dice values between the two sample sizes. Central mark on each violin plot indicates the mean value. fIQ: fluid intelligence. cIQ: crystalized intelligence. IC-Cognition: overall cognitive performance. K-Ridge: kernel ridge regression. CPM: connectome-based predictive modelling. CPM-P, positive associations; CPM-N, negative associations. FDR: false discovery rate. NBS: network-based statistic. ICC: intraclass correlation coefficient.

### Within-sample estimation of feature weight reliability

Previous studies have suggested relatively high feature weight reliability for the predictive models investigated here (Cui and Gong, 2018; Finn and Bandettini, 2021; Finn et al., 2015; Jiang et al., 2019; Rosenberg et al., 2016). Why have we found substantially poorer feature weight reliability? In all the above experiments (Figures 2 and 3), test-retest reliability was evaluated out-of-sample, whereas most previous studies report within-sample estimates based on agreement in beta coefficients across cross-validation folds and iterations. Therefore, we next aimed to explicitly compare within-sample and out-of-sample estimates of prediction accuracy and feature weight test-retest reliability. We focused on ridge regression and compared: i) k-fold (k=10), ii) leave-one-out and iii) half-split (k=2) cross-validation procedures. In the case of k-fold (k=10) and leave-one-out cross-validation, within-sample reliability was estimated by fitting beta coefficients to the pooled sample of k-1 folds and ICC was computed between the resulting k sets of beta coefficients, consistent with common practices in the literature (Cui and Gong, 2018; Finn and Bandettini, 2021; Finn et al., 2015; Jiang et al., 2019; Rosenberg et al., 2016). For out-of-sample reliability, ICC was computed for the averaged feature weights over training sets (Chen et al., 2020; Dhamala et al., 2021; Greene et al., 2018) between the two independent data splits. Leave-one-out and k-fold cross-validation were repeated for 10 genetically unrelated samples of individuals (n=400), given the large amount of computation required, yielding 10 estimates of within-sample ICC (five out-of-sample estimates) and prediction accuracy for each cognitive measure and sex.

We found that correlation coefficients between predicted and actual cognitive performance were highly comparable between the three cross-validation procedures, with comparable variability in accuracy estimates across data samples (Figure 4a). However, within-sample estimates of feature weight test-retest reliability were substantially inflated relative to out-of-sample estimates for all cognitive measures and sex (Figure 4b). Within-sample ICC suggested excellent feature weight reliability (ICC>0.98), whereas out-of-sample estimates suggested poor reliability for cognition and fair reliability for sex prediction. We conclude that within-sample estimates of feature weight reliability can be inflated and out-of-sample estimation should be used, if possible.

**Figure 4.**
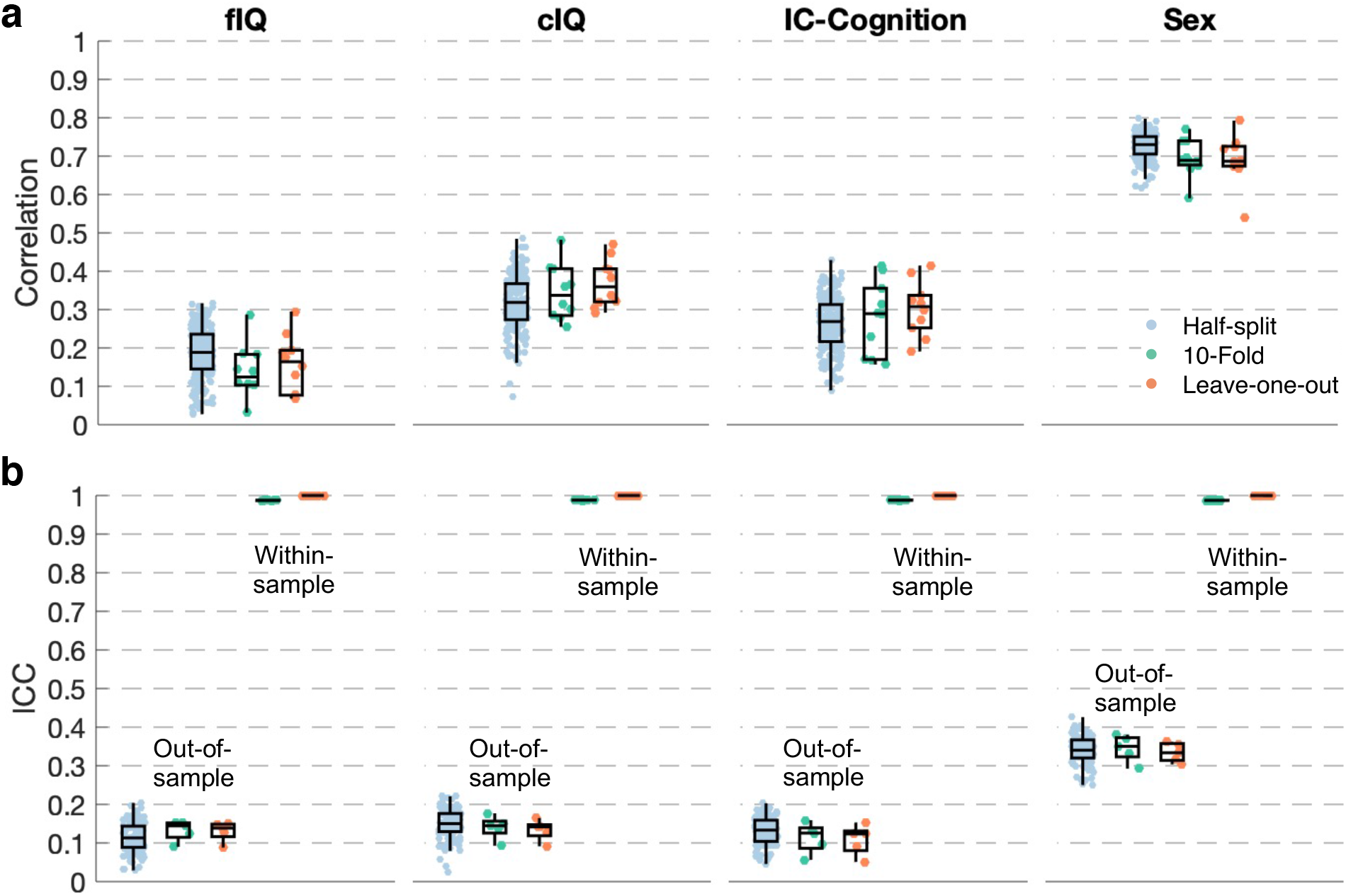
Comparison of within-sample and out-of-sample estimates of feature weight test-retest reliability. **(a)**, Pearson correlation coefficient between observed and out-of-sample predicted fluid intelligence (fIQ), crystalized intelligence (cIQ), overall cognition (IC-Cognition) as well as sex (partial linear correlation, 0-female; 1-male). All predictions are out-of-sample and derived from ridge regression models trained using half-split, 10-fold and leave-one-out cross-validation. Boxplots show sampling variation in correlation coefficients across 100 (half-split) and 10 (10-fold and leave-one-out) random samples of genetically unrelated individuals (n=400). Each data points represents a sample. **(b)**, Out-of-sample and within-sample estimates of feature weight test-retest reliability, quantified with the intraclass correlation coefficient (ICC). Boxplots show sampling variation in ICC across 100 (half-split) and 10 (10-fold and leave-one-out) random samples of genetically unrelated individuals (n=400). The central mark of the box plot indicates the median and the bottom and top edges of the box indicate 25th and 75th percentiles of the distribution, respectively. The whiskers extend to the most extreme data points that are not considered outliers (1.5 × interquartile range).

### Consistency in feature weights between predictive models

We next evaluated the extent of agreement in feature weights between the four predictive models (lasso, ridge and kernel ridge regression and CPM) using ICC. ICC values were computed between all pairs of models using either the same half-split (within-sample ICC) or different half-splits (out-of-sample ICC) for each model in a pair. Figure 5 shows out-of-sample (lower triangle + diagonal) and within-sample mean ICC values (upper triangle), averaged over 100 half-split pairs. This was repeated for n=400 (Figure 5a) and n=800 sample sizes (Figure 5b). We found that feature weights were most consistent between ridge and kernel ridge regression, particularly for sex prediction, but also for the three cognitive measures (within-sample ICC>0.8). Lasso showed fair consistency with the ridge regression (within-sample ICC>0.4), while CPM showed lower consistency with the three regression models for both cognition and sex prediction (within-sample ICC<0.2). The Haufe transformation improved consistency in feature weights between models (Supplementary Figure 2).

**Figure 5.**
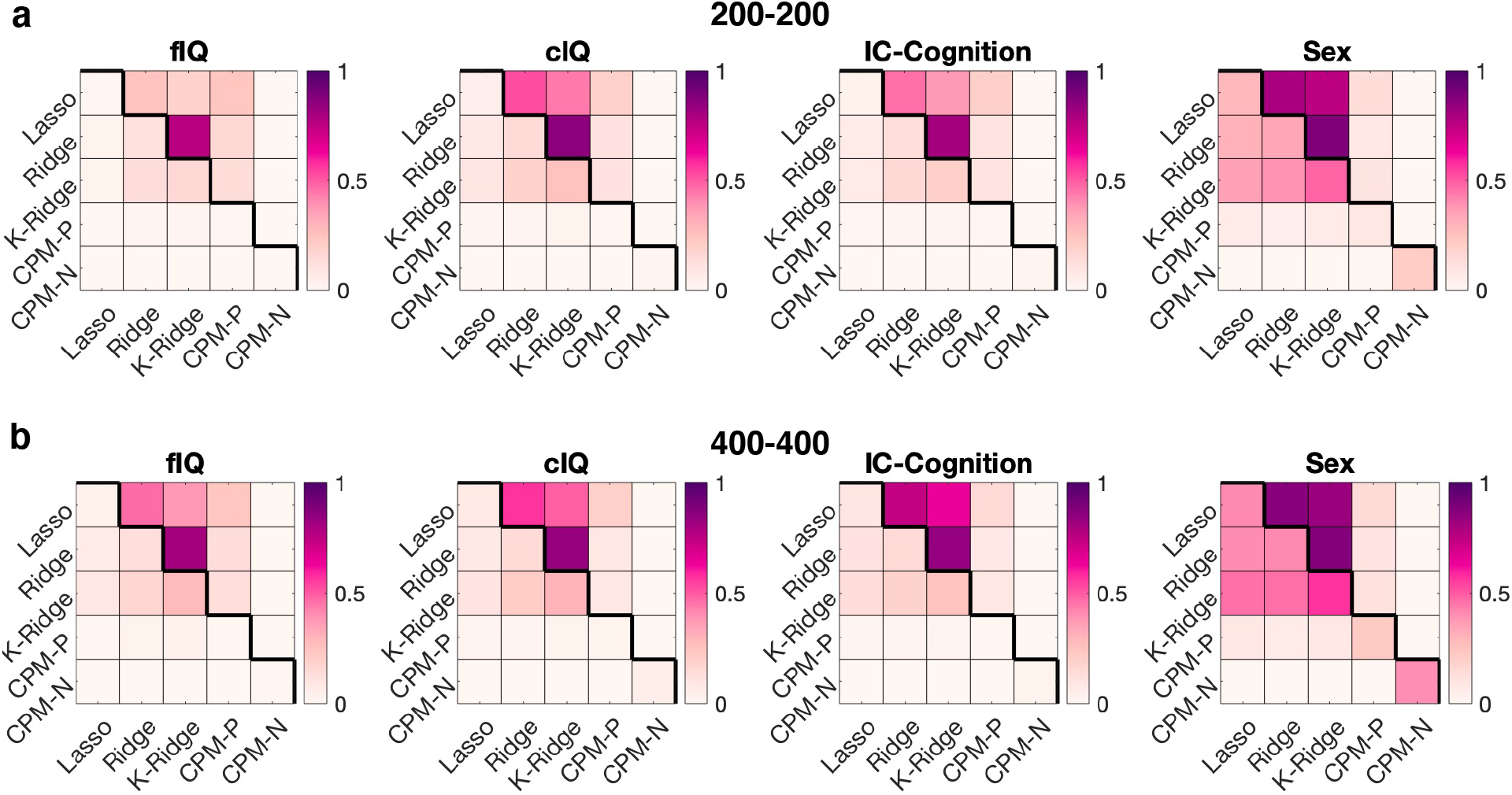
Consistency in feature weights between different predictive models. Matrix cells are colored according to within-sample (upper triangle) and out-of-sample (lower triangle + diagonal) intraclass correlation (ICC) values. ICC values quantify consistency in feature weights estimated by two distinct predictive models and represent averages over 100 half-split pairs. ICC values are shown for samples sizes of n=400, with 200 per half-split **(a)** and n=800, with 400 per half-split **(b)**. fIQ: fluid intelligence. cIQ: crystalized intelligence. IC-Cognition: overall cognitive performance. K-Ridge: kernel ridge regression. CPM: connectome-based predictive modelling. CPM-P, positive associations; CPM-N, negative associations.

### Regional analysis

In all the above experiments, feature weight test-retest reliability was quantified globally, without regard for possible differences in reliability between connections and regions comprising the feature space. Therefore, we next investigated regional variation in feature weight reliability in the case of ridge regression prediction of fIQ. We considered the symmetric 376 × 376 matrix of feature weights, where element (*i, j*) is the estimated beta coefficient for the connection between regions *i* and *j*. Summing across either the positive or negatives values in the rows of this matrix provided a region-specific characterization of feature importance, which we visualized on the cortical surface for Split 1 and Split 2 of a representative half-split pair. These regional features weights can be interpreted such that higher fluid intelligence is predicted for individuals with stronger connectivity with positively weighted regions and lower (negative) connectivity with negatively weighted regions. We observed substantial regional variation in summed feature weights between Split 1 and Split 2 (Figure 6a). The Haufe transformation marginally reduced variation between half-split pairs, although reliable attribution of feature importance to specific regions remained challenging (Figure 6b). For example, the temporal cortex was associated with positive feature weights in Split 1, whereas the prefrontal cortex and striatum were weighted more positively in Split 2 (Figure 6b). Similar variation was evident for negatively weighted regions, where the ventromedial prefrontal cortex was weighted most negatively in Split 1, but regions of the parietal cortex and temporal pole were instead associated with negative feature weights in Split 2 (Figure 6b). In contrast, feature importance could be ascribed more reliably to specific regions and canonical brain networks in the case of sex prediction (Figure 6c & 6d). For example, the most prominent predictive features of male sex were strong between-network connections, particularly between regions in the default mode network and regions in other networks, such as the dorsal attention, visual and ventral attention network. Contrastingly, strong within-network connections were generally more predictive of females. Following the Haufe transformation, male-predictive features showed significantly greater spatial consistency on average, compared to female-predictive features (median±SD across 100 half-split pairs: male: r=0.53±0.29; female: r=0.38±0.35; t=3.87, p=1.5×10^−4^); however, sex differences were not observed in the original feature space (male: r=0.29±0.06; female: r=0.31±0.60; t=1.38, p=0.17). Taken together, these regional analyses suggest that reliably explaining predictive utility in terms of specific brain regions and canonical functional networks is challenging for predictive models of cognitive performance.

**Figure 6.**
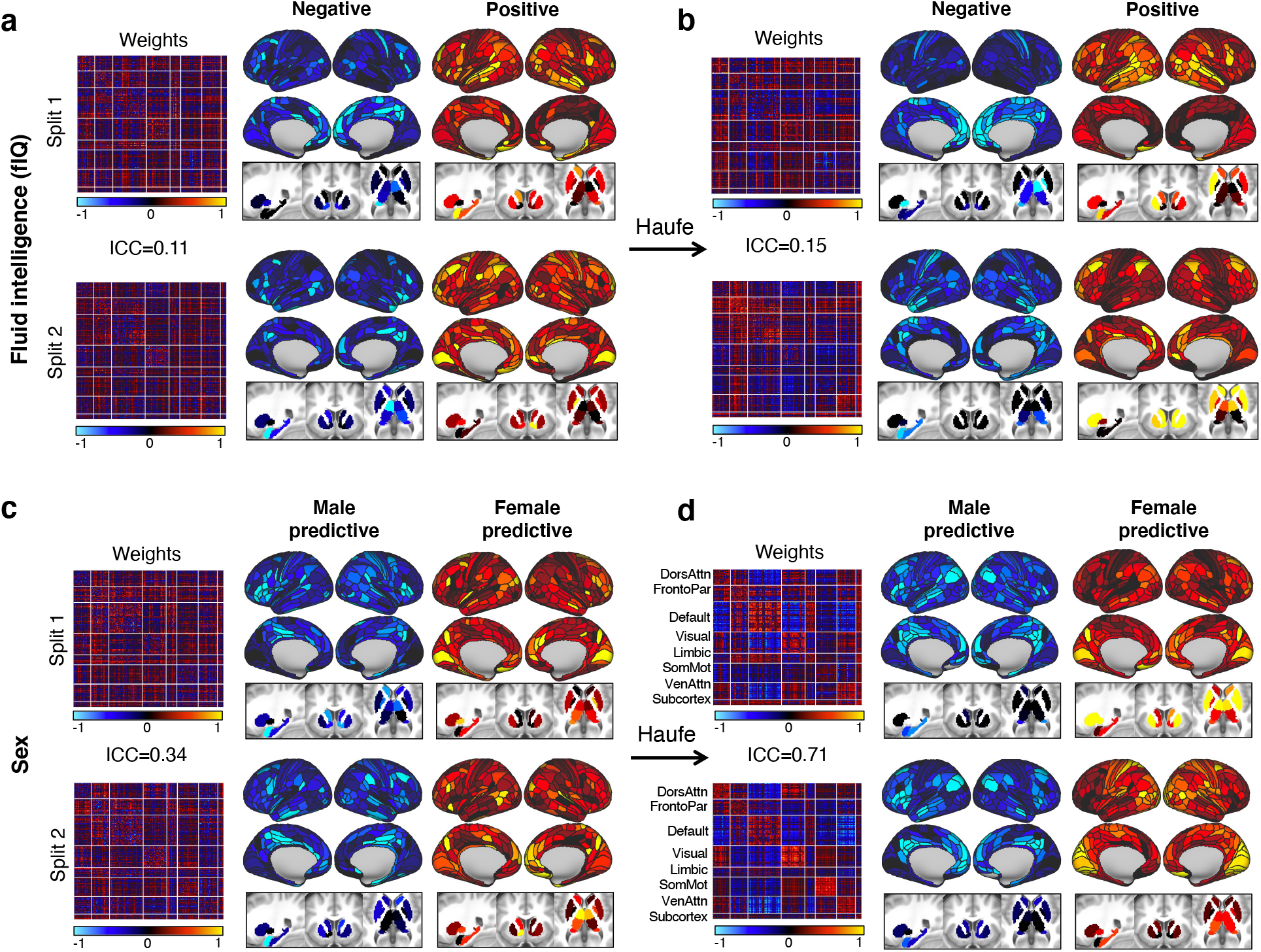
Regional representation of connectivity feature weights. Estimated connectivity feature weights are shown as a 376 × 376 matrix, where element 0*i, j*) is the estimated beta coefficient for the connection between regions *i* and *j*. Matrices are shown for Split 1 and Split 2 of a representative half-split pair (i.e., the half-split pair closest to the median intraclass correlation coefficient, ICC). Matrix rows/columns are ordered to group regions according to large-scale functional brain networks, as demarcated by solid white lines (Yeo et al., 2011). Positive and negative feature weights were summed separately across each matrix row to provide a regional characterization of feature importance. Cortical renderings and subcortical slices of regional feature weights are shown for ridge regression prediction of fluid intelligence **(a, b)** and sex **(c, d)**. Feature weights are shown with **(b, d)** and without **(a, c)** the Haufe transformation. Higher fluid intelligence (or female sex) is predicted for individuals with stronger connectivity with positively weighted regions and lower (negative) connectivity with negatively weighted regions. DorsAttn, dorsal attention network; FrontoPar, frontoparietal network; SmoMot, somatomotor network; VenAttn, ventral attention network.

### Impact of feature space dimensionality

Finally, we investigated the impact of feature space dimensionality on prediction accuracy and feature weight rest-retest reliability. The above experiments were based on a relatively high-dimensional feature space comprising 70,500 connections (376 nodes). We therefore reduced feature space dimensionality by using progressively coarser whole-brain parcellation atlases previously mapped with spatial independent component analysis (ICA; see Methods). We considered atlases comprising 15, 25, 50, 100, 200 and 300 spatial ICA components (nodes), yielding 105, 300, 1225, 4950, 19900 and 44850 connectivity features, respectively. For each dimensionality, prediction accuracy and feature weight test-retest reliability were evaluated using ridge regression and the same half-split procedure described above.

We found that correlation coefficients between predicted and actual cognitive performance increased with feature space dimensionality, particularly for sex prediction, and to a lesser extent for fIQ, cIQ and IC-Cognition (Figure 7a). However, gains in prediction accuracy achieved through increased feature space dimensionality were at the expense of poorer feature weight test-retest reliability (Figure 7b). Specifically, for all three cognitive measures, mean ICC values increased from less than 0.1 for the highest dimensional feature space (300 ICA components), to above 0.3 for the lowest dimensional feature space (15 components). Therefore, a more than two-fold increase in ICC can be achieved by reducing feature space dimensionality from 300 to 15 nodes, albeit at the cost of a 9-45% reduction in prediction accuracy. Interestingly, the cortex (Glasser et al., 2016a) and subcortex (Tian et al., 2020) atlases (376 nodes) provided a feature space that was more reliable than the comparably sized feature space derived from spatial ICA components (300 nodes). We conclude that the choice of brain parcellation and resolution at which functional connectivity is mapped provides a means to arbitrate an inherent tradeoff between prediction accuracy and feature weight reliability.

**Figure 7.**
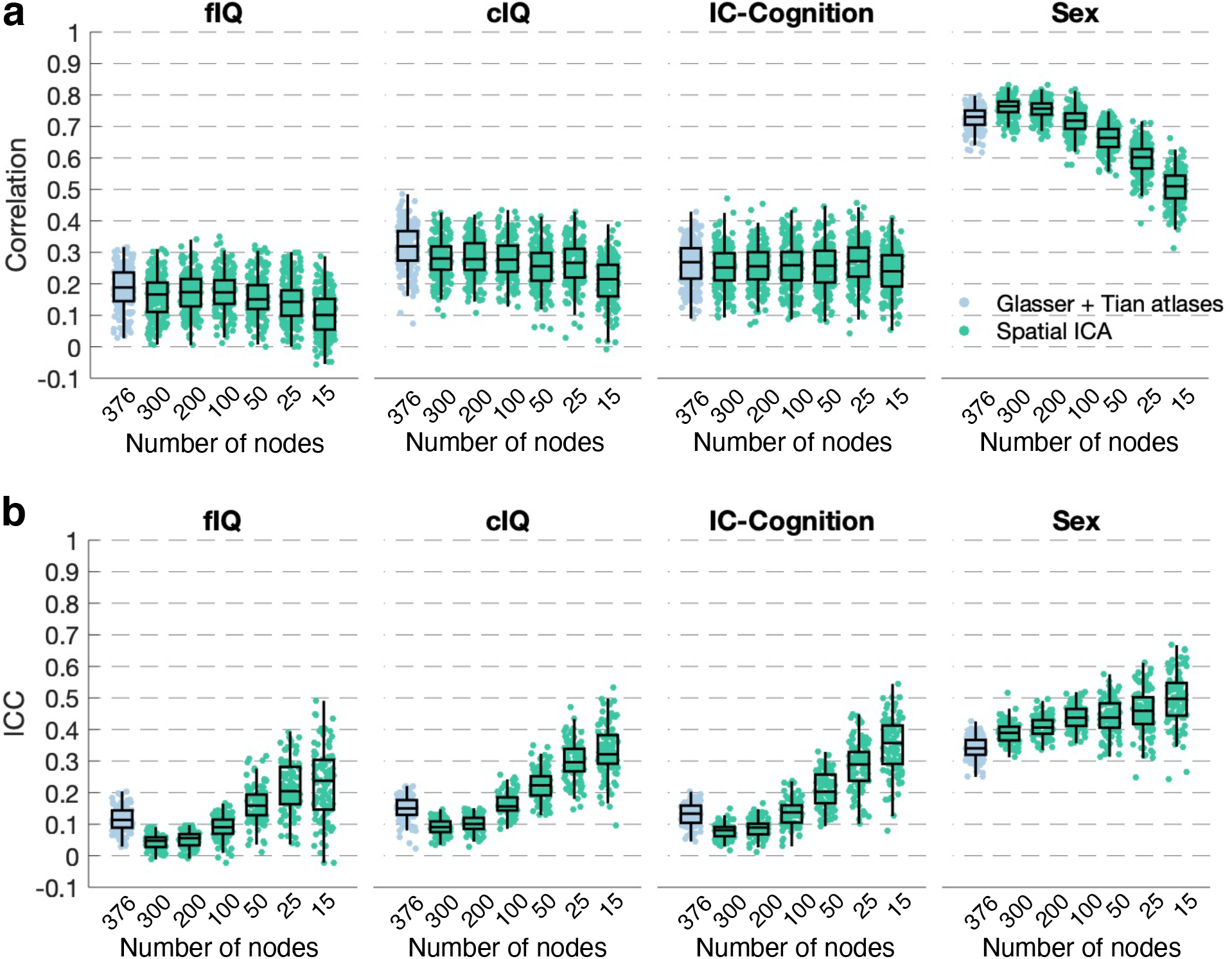
Tradeoff between prediction accuracy and feature weight test-retest reliability arbitrated by feature space dimensionality. **(a)**, Pearson correlation coefficients between observed and out-of-sample predicted fluid intelligence (fIQ), crystalized intelligence (cIQ), overall cognition (IC-Cognition) as well as sex (partial linear correlation, 0-female; 1-male). Boxplots characterize variation in correlation coefficients across 100 half-split samples and are shown for six feature space dimensionalities derived from spatial ICA components (number of nodes: 15, 25, 50, 100, 200, 300; green). Boxplots are also shown for connectivity features derived from established parcellation atlases (376 nodes, blue). All predictions computed using ridge regression. **(b)**, Feature weight test-retest reliability (out-of-sample) quantified with the intraclass correlation coefficient (ICC). Boxplots characterize variation in ICC across 100 half-split samples and are shown for the same feature space dimensionalities. The central mark of the box plot indicates the median and the bottom and top edges of the box indicate 25th and 75th percentiles of the distribution, respectively. The whiskers extend to the most extreme data points that are not considered outliers (1.5 × interquartile range).

## Discussion

Although cognitive performance and intelligence can be reliably predicted with modest accuracy from an individual’s resting-state functional connectivity, we found that feature weight test-retest reliability is poor. Reliably mapping predictive importance to specific connections, regions and networks is therefore challenging. Poor feature weight reliability limits the extent to which a machine learning approach can be used to explain neurobiological mechanisms of cognition and to test theoretical cognitive models. We found that large sample sizes, certain feature weight transformations and connectivity mapped at coarse spatial resolutions marginally improved feature weight reliability. However, ICC values between pairs of feature weights remained poor (ICC<0.4), even in the most favorable settings. In contrast, using the same features to predict an individual’s sex yielded substantially more reliable feature weights, suggesting that the integrity of the features themselves (i.e., resting-state functional connectivity measurements) was not entirely to blame for poor feature weight reliability when predicting cognitive performance.

While providing a barrier to explainable machine learning, *not* being able to reliably localize predictive utility to specific connectivity features can provide clues about the neural basis of cognition. Poor feature weight reliability may be a consequence of the *dynamic* nature of resting-state functional connectivity and significant heterogeneity among individuals in preferred cognitive strategies to make decisions (Finn and Rosenberg, 2021; Marewski and Schooler, 2011). Given that different cognitive strategies can differentially impact functional connectivity (Park et al., 2019), in a diverse sample of individuals, there may be multiple feature weight solutions that achieve comparable prediction accuracy, each of which characterizes a distinct cognitive strategy. In this scenario, a machine learning algorithm will learn only one of the many solutions, without providing explicit insight into alternative solutions providing comparable prediction accuracy. A change in the composition of the training sample through addition of individuals favoring a certain cognitive strategy could, in principle, result in the model learning an alternative solution. Therefore, given that cognitive functions are complex and heterogenous, and resting-state functional connectivity is an inherently dynamic phenotype, we hypothesize that broad measures of cognitive performance, such as fluid and crystalized intelligence, cannot be neatly pinned down to a unique set of connectivity features.

This hypothesis is supported by the limited reproducibility among classical significance testing studies that have sought to map group-level associations between complex behavioral traits and resting-state functional connectivity (Poldrack et al., 2017). If distinct functional circuits are indeed engaged by different cognitive strategies, group-level network maps will likely capture a mix of circuits inherent to each strategy, and thus might not necessarily represent any given individual comprising the sample.

Throughout this study, the extent of agreement in feature weights between half-split samples was interpreted from the standpoint of test-retest reliability. However, low ICC values between feature weights can also be construed as evidence for feature selection instability and solution non-uniqueness (machine learning perspective), sampling variability (statistical perspective) and poor measurement validity (cognitive psychology perspective). Instability in feature selection is a well-known issue in machine learning, where small changes to the training sample can lead to large changes in feature weights (Nogueira et al., 2017). However, we confirmed that stochasticity in hyperparameter optimization and the model fitting process introduced minimal instability. Regarding measurement validity, the instruments used to measure fluid and crystallized intelligence in the Human Connectome Project are validated and extensively used (Bilker et al., 2012; Weintraub et al., 2013). For these reasons, we believe that our findings are most appropriately contextualized in terms of test-retest reliability, although all three of the above perspectives are relevant and interrelated.

Several recent studies have sought to predict cognitive performance from an individual’s resting-state functional connectivity. Most recently, Dhamala and colleagues found that distinct functional and structural connections predict fluid and crystallized intelligence (Dhamala et al., 2021). In earlier work, Kong and colleagues considered predicting cognitive performance from individual-specific functional brain networks. They found that individualized cortical topographies generalized more readily to new fMRI data from the same individual, compared to group-consensus parcellations (Kong et al., 2019). Li and colleagues found that global signal regression—a controversial fMRI preprocessing step—improved prediction accuracy (Li et al., 2019). Several recent studies have focused on comparing deep and machine learning approaches for neuroimaging-based prediction of cognitive performance and behavioral traits (Abrol et al., 2021; He et al., 2020; Schulz et al., 2020). While evaluating and maximizing prediction accuracy is a key consideration of these previous studies, some previous studies have also evaluated consensus in feature importance for the purpose of visualization or external validation of the accuracy of predictive models (Cui and Gong, 2018; Finn and Bandettini, 2021; Finn et al., 2015; Jiang et al., 2019; Rosenberg et al., 2016). To the best of our knowledge, while these previous studies did not explicitly evaluate feature weight reliability, consensus in feature importance was computed based on agreement in feature weights between cross-validation folds or iterations. Several of these studies observed relatively high consensus in feature importance between cross-validation folds, but they did not necessarily claim high feature weight reliability based on this observation. We found that such within-sample measurements can lead to inflated estimates of feature weight reliability (Figure 4). Out-of-sample estimates computed using rigorous half-split experiments were found to be substantially lower than previous studies suggested and suggest poor feature weight test-retest reliability.

We identified a tradeoff between prediction accuracy and feature weight reliability. Specifically, increasing feature space dimensionality by using higher resolution parcellation atlases led to increased prediction accuracy, at the expense of poorer feature weight reliability. Improved reliability could be due to the higher signal-to-noise ratio afforded by averaging fMRI signals over broader spatial extents, leading to more accurate functional connectivity measurements, compared to those derived from high-resolution parcellation atlases. Reliable interpretation of feature importance can therefore benefit from low-dimensional connectome mapping, if the decrease in prediction accuracy and spatial resolution can be tolerated.

Some remarks on the relative performance of the four predictive models are warranted. Feature weight reliability was poor for all predictive models and cognitive measures, although subtle differences were evident. Kernel ridge and ridge regression yielded the most reliable feature weights and the most accurate predictions, whereas the performance of lasso and CPM was lower for both measures. Unlike the other three predictive models, CPM uses a univariate feature selection strategy, and this may impact feature weight reliability. O’Connor and colleagues recently incorporated resample aggregation into CPM, which was found to improve CPM generalizability (O’Connor et al. 2021). This resampling procedure can provide insight into feature weight reliability and stability. The authors report that fewer than 1% of the 35,778 connectivity features were selected in more than 90% of the resamples (200-subsample: 1.95 ± 1.5 connections; 300-subsample: 58 ± 20; bagged model: 46 ± 16). While these percentages are not directly comparable with the Dice coefficients reported here, they also suggest considerable variation in selected features between samples. Another difference obscuring direct comparison is that we compared feature selection between independent samples of individuals, whereas individuals can be included in more than one of the samples generated by O’Connor and colleagues. We also note that CPM prediction accuracies for fIQ reported by these authors are marginally higher than those reported here. This may be due to the use of different parcellation atlases, samples and cross-validation to optimize the *p*-value feature selection threshold. CPM is an important predictive model and further work is needed to evaluate feature selection reliability under resample aggregation.

We also investigated the test-retest reliability of mass univariate significance testing. For each connection, this involved independently testing the null hypothesis of an absence of association between cognitive performance and resting-state functional connectivity strength. Interestingly, the connection-wise univariate test statistic used to assess this null hypothesis showed greater reliability than predictive feature weights. Therefore, if the researcher’s principal goal is to elucidate relationships between cognitive performance and brain connectivity, classical statistical inference should not be overlooked because it can provide improved reliability compared to predictive modelling. However, the set of significant connections derived from thresholding the corresponding *p*-values to control the family-wise error or false discovery rate showed low Dice coefficients between half-split pairs. We conclude that while dichotomization based on statistical significance can aid interpretation by categorically localizing effects to distinct connections, continuous measures of association between cognitive performance and connectivity strength are more reliable.

Using larger samples may improve feature weight reliability, although we found the doubling the sample size from n=400 to n=800 led to only a marginal increase in ICC values. Increasing the sample size beyond n=800 may gradually saturate the reliability estimates or approach a plateau (Schulz et al., 2020). However, studies often only focus on the impact of sample sizes on prediction accuracies (Marek et al., 2020; Poldrack et al., 2020; Schulz et al., 2020; Varoquaux, 2018) and feature weight test-retest reliability as a function of sample size remains to be characterized using even larger neuroimaging datasets such as the UK Biobank (Miller et al., 2016). Future work should also focus on investigating the reliability of predictive models derived from task-evoked fMRI, given that task-evoked brain connectivity can yield more accurate predictions of cognitive performance compared to resting-state connectivity (Chen et al., 2020; Finn and Bandettini, 2021; Greene et al., 2018; Jiang et al., 2020). Future studies should also investigate the impact of personalized brain atlases (Fair, 2018; Gordon et al., 2017; Kong et al., 2019; Wang et al., 2020) on the reliability of feature weights. Additionally, future evaluations incorporating multiple independent datasets would enable testing of the generalizability of feature weights between different cohorts acquired at different sites. This is a stronger form of out-of-sample test-retest reliability than the one evaluated here. For example, Rosenberg and colleagues (2016; 2018b) identified networks predicting attentional task performance in two independent datasets and report statistically significant overlap between the two networks, providing evidence of model reliability and generalizability. This measure of overlap is not necessarily comparable to the feature weight reliability reported here based on the ICC.

Finally, it is important to remark that feature importance can be deduced in ways other than interrogating feature weights. Certain features can be excluded from the feature space and the predictive models retrained using the reduced feature space. Any reduction in prediction accuracy provides an indirect measure of the importance of the omitted features (Yarkoni and Westfall, 2017). While this approach has not been widely adopted for functional connectivity, perhaps due to high feature space dimensionality, Cropley and colleagues systematically omitted specific brain lobes from a predictive model of brain age based on gray matter morphology. They found that omitting frontal regions reduced the strength of the association between brain age gap and psychosis symptoms, providing evidence for the importance of the frontal lobes in psychopathology (Cropley et al., 2021). The reliability of feature importance estimates deduced from this hierarchical model comparison approach remains to be investigated.

### Conclusion

Neuroimaging-based predictive models in cognitive neuroscience are burgeoning. However, predicting for the sake of prediction is an easy trap to fall into and we hope that our work prompts a rebalance of attention from maximizing prediction accuracy to establishing explainable models built on reliable feature weights. For current predictive models of cognitive performance based on resting-state functional connectivity, feature importance is difficult to reliably estimate, meaning that localizing predictive utility to specific connections and circuits is challenging. This limits the extent to which predictive models can be explained in terms of neurobiological mechanisms. We found that larger sample sizes, coarser parcellation atlases and non-sparse feature selection/regularization can marginally improve feature weight test-retest reliability. We recommend estimating reliability out-of-sample, if possible. The more common approach of measuring agreement in feature weights between cross-validation folds and iterations provides inflated estimates of feature weight reliability.

## Data and code availability statement

All data analyzed in this study are available for download to anyone agreeing to the Open Access Data Use Terms (https://db.humanconnectome.org/). Access to family structure data and several behavioral measures requires acceptance of the HCP Restricted Data Use Terms (https://www.humanconnectome.org/study/hcp-young-adult/document/restricted-data-usage). Matlab code for conducting the core analyses is available on GitHub (https://github.com/yetianmed/MLReliability).

## Credit authorship contribution statement

**Ye Tian:** Conceptualization, data curation, software, analysis, visualization, writing.

**Andrew Zalesky:** Conceptualization, analysis, writing, funding, supervision.

## Acknowledgments

Data were provided by the Human Connectome Project, WU-Minn Consortium (Principal Investigators: David Van Essen and Kamil Ugurbil; 1U54MH091657) funded by the 16 NIH Institutes and Centers that support the NIH Blueprint for Neuroscience Research; and by the McDonnell Center for Systems Neuroscience at Washington University. AZ is supported by the NHMRC Senior Research Fellowship (APP1142801).

## Supplementary Materials

### Regularization and hyperparameter optimization

A nested 20-fold cross-validation was used for hyperparameter optimization for the three regression methods, including lasso (Tibshirani 1996), ridge (Hoerl and Kennard 1970) and kernel ridge regression (Li et al. 2019). This was performed using the MATLAB function *fitrlinear*.*m* for continuous cognitive measures and *fitclinear*.*m* for the binary sex variable. The nested 20-fold cross-validation was performed for each half-split cross-validation iteration. For each inner training iteration, a grid search was used to randomly search over a sequence of 100 logarithmically spaced hyperparameters (*λ*), ranging from 10^−5^/N to 10^5^/N, where N is the number of training subjects comprising the inner fold. The sparse reconstruction by separable approximation (SpaRSA) technique (Wright et al. 2009) was used to minimize the objective function for lasso, and a combination of average stochastic gradient descent (ASGD) and limited-memory Broyden-Fletcher-Goldfarb-Shanno quasi-Newton algorithm (LBFGS) (Nocedal and Wright 2006) was used for ridge and kernel ridge regression. A maximal number of 1000 iterations were specified, and the optimization terminated when 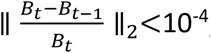, where *B*_*t*_ is the regression coefficients and the intercept at optimization iteration *t*. The *λ* value resulting in the smallest error across all inner test folds were selected to compute the beta coefficients using all individuals in the outer training split.

Similarly, a nested 20-fold cross-validation was used for the connectome-based predictive modeling (CPM) (Shen et al. 2017), in which a sequence of 100 logarithmically spaced *p*-values ranging from 10^−5^ to 1 was tested. The *p*-value threshold resulting in maximal positive or negative association between the summation of functional connectivity (FC) strengths and cognitive performance (or sex) was selected to compute the summation of FC strengths across all individuals in the outer training split.

**Figure S1.**
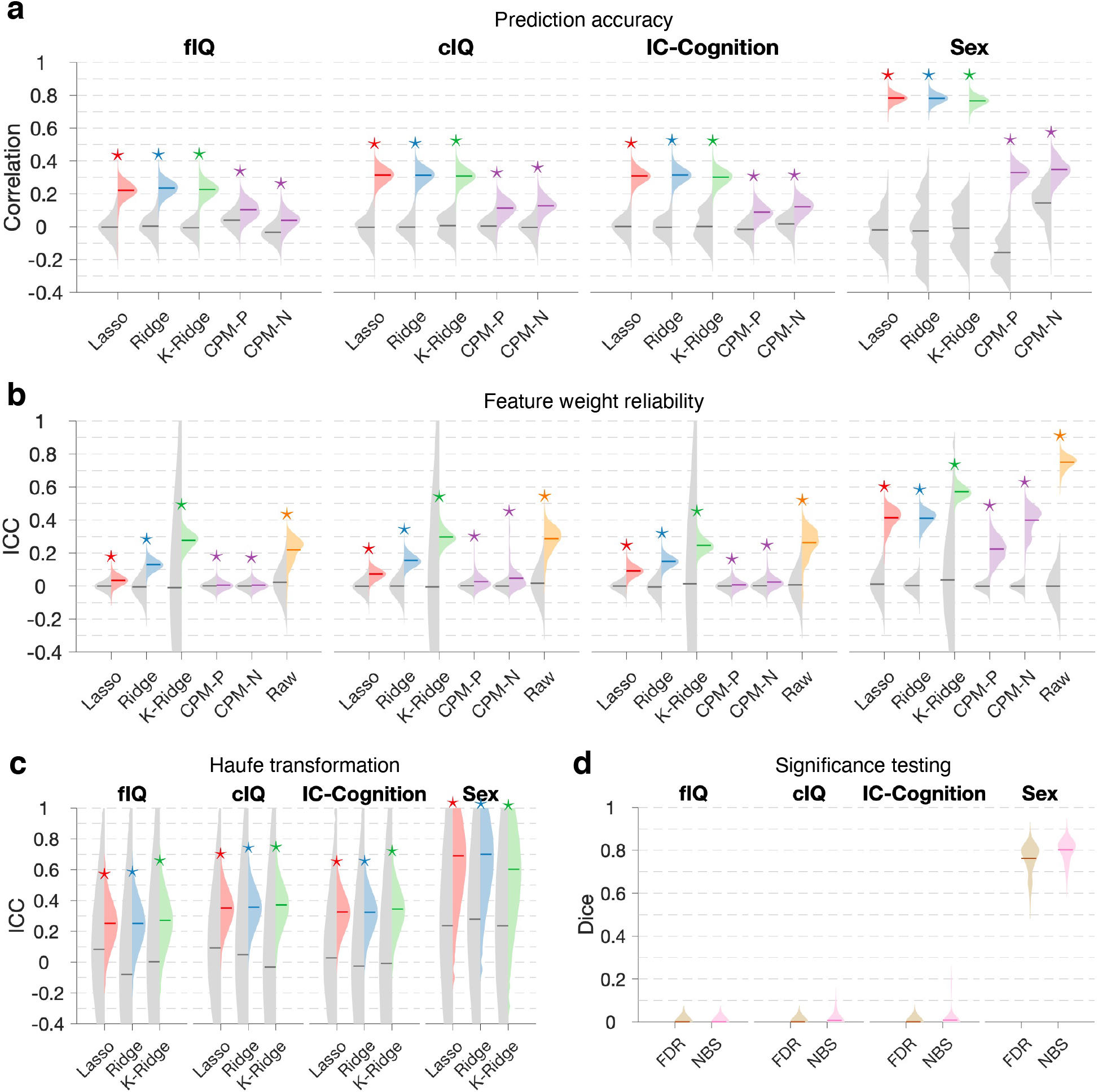
Prediction accuracy and feature weight test-retest reliability estimated using half-split cross-validation in 800 individuals. **(a)**, Pearson correlation coefficient between observed and out-of-sample predicted fluid intelligence (fIQ), crystalized intelligence (cIQ), overall cognition (IC-Cognition) as well as sex (partial linear correlation, 0-female; 1-male) using Lasso regression, ridge regression, kernel ridge regression and connectome-based predictive modeling (CPM). CPM-P, positive associations; CPM-N, negative associations. Violin plots show the distribution of correlation coefficients across 100 half-split pairs (left: chance; right: observed). Asterisks are shown to indicate mean Pearson correlation coefficients significantly exceeding chance-level predictions (two-sample t-test, p<0.05, false discovery rate (FDR) correction across five models). **(b)**, Feature weight test-retest reliability (out-of-sample) quantified with intraclass correlation coefficient (ICC) for the five predictive models and the raw test statistic. Violin plots show the distribution of ICC across 100 half-split pairs (left: chance; right: observed). Asterisks are shown to indicate ICC mean values significantly greater than chance (two-sample t-test, p<0.05, FDR correction across six models). Raw refers to the univariate test statistic for each connection assessing the null hypothesis of an absence of association between cognitive performance and functional connectivity strength. **(c)**, Same as (b), but with the Haufe transformation applied to feature weights before evaluating test-retest reliability. **(d)**, The network-based statistic (NBS) and false discovery rate (FDR) were used to identify connections with functional connectivity strengths that significantly correlated with cognitive performance (family-wise error rate and FDR controlled at 0.05, respectively). Violin plots show distribution of dice coefficients quantifying overlap in connections declared statistically significant between 100 half-split pairs. Central mark on each violin plot indicates the mean value.

**Figure S2.**
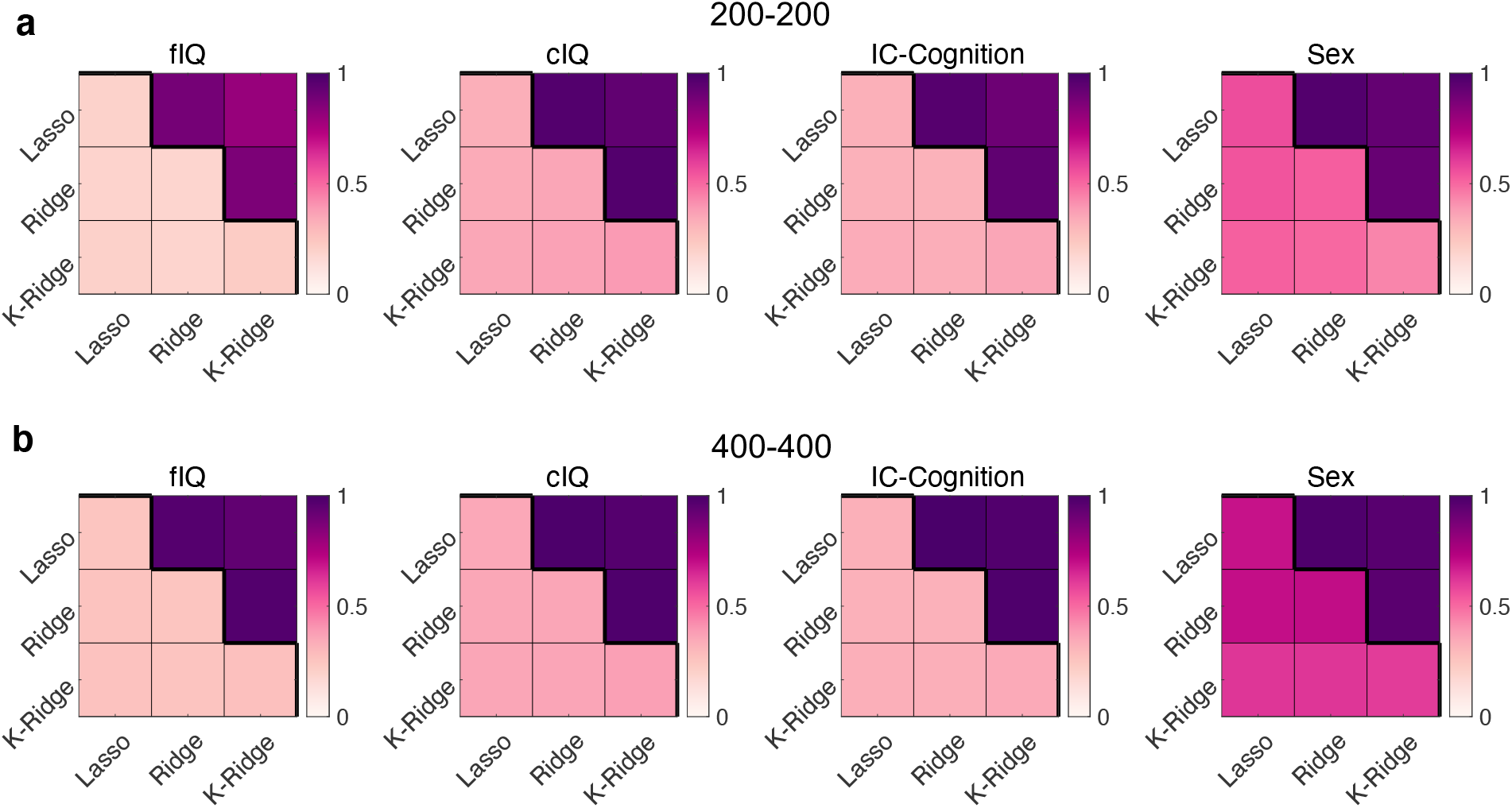
Consistency in feature weights between different predictive models with the Haufe transformation. Matrix cells are colored according to within-sample (upper triangle) and out-of-sample (lower triangle + diagonal) intraclass correlation (ICC) values. ICC values quantify consistency in feature weights with the Haufe transformation estimated by two distinct predictive models and represent averages over 100 half-split pairs. ICC values are shown for samples sizes of n=400, with 200 per half-split **(a)** and n=800, with 400 per half-split **(b)**. fIQ: fluid intelligence. cIQ: crystalized intelligence. IC-Cognition: overall cognitive performance. K-Ridge: kernel ridge regression. CPM: connectome-based predictive modelling. CPM-P, positive associations; CPM-N, negative associations.

## References

Abrol, A., Fu, Z., Salman, M., Silva, R., Du, Y., Plis, S., Calhoun, V., 2021. Deep learning encodes robust discriminative neuroimaging representations to outperform standard machine learning. Nature Communications 12, 353.

Beckmann, C.F., Smith, S.M., 2004. Probabilistic independent component analysis for functional magnetic resonance imaging. IEEE transactions on medical imaging 23, 137–152.

Bilker, W.B., Hansen, J.A., Brensinger, C.M., Richard, J., Gur, R.E., Gur, R.C., 2012. Development of abbreviated nine-item forms of the Raven’s standard progressive matrices test. Assessment 19, 354–369.

Birn, R.M., Molloy, E.K., Patriat, R., Parker, T., Meier, T.B., Kirk, G.R., Nair, V.A., Meyerand, M.E., Prabhakaran, V., 2013. The effect of scan length on the reliability of resting-state fMRI connectivity estimates. NeuroImage 83, 550–558.

Bzdok, D., Engemann, D., Thirion, B., 2020. Inference and Prediction Diverge in Biomedicine. Patterns (N Y) 1, 100119.

Chen, J., Tam, A., Kebets, V., Orban, C., Ooi, L.Q.R., Marek, S., Dosenbach, N., Eickhoff, S., Bzdok, D., Holmes, A.J., Thomas Yeo, B.T., 2020. Shared and unique brain network features predict cognition, personality and mental health in childhood. bioRxiv, 2020.2006.2024.168724.

Cropley, V.L., Tian, Y., Fernando, K., Mansour L S., Pantelis, C., Cocchi, L., Zalesky, A., 2021. Brain-Predicted Age Associates With Psychopathology Dimensions in Youths. Biological Psychiatry: Cognitive Neuroscience and Neuroimaging 6, 410–419.

Cui, Z., Gong, G., 2018. The effect of machine learning regression algorithms and sample size on individualized behavioral prediction with functional connectivity features. NeuroImage 178, 622–637.

Dhamala, E., Jamison, K.W., Jaywant, A., Dennis, S., Kuceyeski, A., 2021. Distinct functional and structural connections predict crystallised and fluid cognition in healthy adults. Hum Brain Mapp.

Eickhoff, S.B., Langner, R., 2019. Neuroimaging-based prediction of mental traits: Road to utopia or Orwell? PLOS Biology 17, e3000497.

Fair, D.A., 2018. The Big Reveal: Precision Mapping Shines a Gigantic Floodlight on the Cerebellum. Neuron 100, 773–776.

Filippini, N., MacIntosh, B.J., Hough, M.G., Goodwin, G.M., Frisoni, G.B., Smith, S.M., Matthews, P.M., Beckmann, C.F., Mackay, C.E., 2009. Distinct patterns of brain activity in young carriers of the APOE-ε4 allele. Proceedings of the National Academy of Sciences 106, 7209–7214.

Finn, E.S., Bandettini, P.A., 2021. Movie-watching outperforms rest for functional connectivity-based prediction of behavior. NeuroImage 235, 117963.

Finn, E.S., Rosenberg, M.D., 2021. Beyond fingerprinting: Choosing predictive connectomes over reliable connectomes. NeuroImage 239, 118254.

Finn, E.S., Shen, X., Scheinost, D., Rosenberg, M.D., Huang, J., Chun, M.M., Papademetris, X., Constable, R.T., 2015. Functional connectome fingerprinting: identifying individuals using patterns of brain connectivity. Nat Neurosci 18, 1664.

Glasser, M.F., Coalson, T.S., Bijsterbosch, J.D., Harrison, S.J., Harms, M.P., Anticevic, A., Van Essen, D.C., Smith, S.M., 2018. Using temporal ICA to selectively remove global noise while preserving global signal in functional MRI data. NeuroImage 181, 692–717.

Glasser, M.F., Coalson, T.S., Robinson, E.C., Hacker, C.D., Harwell, J., Yacoub, E., Ugurbil, K., Andersson, J., Beckmann, C.F., Jenkinson, M., Smith, S.M., Van Essen, D.C., 2016a. A multi-modal parcellation of human cerebral cortex. Nature 536, 171–178.

Glasser, M.F., Smith, S.M., Marcus, D.S., Andersson, J.L., Auerbach, E.J., Behrens, T.E., Coalson, T.S., Harms, M.P., Jenkinson, M., Moeller, S., Robinson, E.C., Sotiropoulos, S.N., Xu, J., Yacoub, E., Ugurbil, K., Van Essen, D.C., 2016b. The Human Connectome Project’s neuroimaging approach. Nat Neurosci 19, 1175–1187.

Glasser, M.F., Sotiropoulos, S.N., Wilson, J.A., Coalson, T.S., Fischl, B., Andersson, J.L., Xu, J., Jbabdi, S., Webster, M., Polimeni, J.R., Van Essen, D.C., Jenkinson, M., 2013. The minimal preprocessing pipelines for the Human Connectome Project. NeuroImage 80, 105–124.

Gordon, E.M., Laumann, T.O., Gilmore, A.W., Newbold, D.J., Greene, D.J., Berg, J.J., Ortega, M., Hoyt-Drazen, C., Gratton, C., Sun, H., Hampton, J.M., Coalson, R.S., Nguyen, A.L., McDermott, K.B., Shimony, J.S., Snyder, A.Z., Schlaggar, B.L., Petersen, S.E., Nelson, S.M., Dosenbach, N.U.F., 2017. Precision Functional Mapping of Individual Human Brains. Neuron 95, 791-807.e797.

Gratton, C., Laumann, T.O., Nielsen, A.N., Greene, D.J., Gordon, E.M., Gilmore, A.W., Nelson, S.M., Coalson, R.S., Snyder, A.Z., Schlaggar, B.L., Dosenbach, N.U.F., Petersen, S.E., 2018. Functional Brain Networks Are Dominated by Stable Group and Individual Factors, Not Cognitive or Daily Variation. Neuron 98, 439-452.e435.

Greene, A.S., Gao, S., Scheinost, D., Constable, R.T., 2018. Task-induced brain state manipulation improves prediction of individual traits. Nature Communications 9, 2807.

Griffanti, L., Salimi-Khorshidi, G., Beckmann, C.F., Auerbach, E.J., Douaud, G., Sexton, C.E., Zsoldos, E., Ebmeier, K.P., Filippini, N., Mackay, C.E., Moeller, S., Xu, J., Yacoub, E., Baselli, G., Ugurbil, K., Miller, K.L., Smith, S.M., 2014. ICA-based artefact removal and accelerated fMRI acquisition for improved resting state network imaging. NeuroImage 95, 232–247.

Haufe, S., Meinecke, F., Görgen, K., Dähne, S., Haynes, J.-D., Blankertz, B., Bießmann, F., 2014. On the interpretation of weight vectors of linear models in multivariate neuroimaging. NeuroImage 87, 96–110.

He, T., Kong, R., Holmes, A.J., Nguyen, M., Sabuncu, M.R., Eickhoff, S.B., Bzdok, D., Feng, J., Yeo, B.T.T., 2020. Deep neural networks and kernel regression achieve comparable accuracies for functional connectivity prediction of behavior and demographics. NeuroImage 206, 116276.

Hoerl, A.E., Kennard, R.W., 1970. Ridge Regression: Biased Estimation for Nonorthogonal Problems. Technometrics 12, 55–67.

Jiang, R., Calhoun, V.D., Fan, L., Zuo, N., Jung, R., Qi, S., Lin, D., Li, J., Zhuo, C., Song, M., Fu, Z., Jiang, T., Sui, J., 2019. Gender Differences in Connectome-based Predictions of Individualized Intelligence Quotient and Sub-domain Scores. Cerebral Cortex 30, 888–900.

Jiang, R., Zuo, N., Ford, J.M., Qi, S., Zhi, D., Zhuo, C., Xu, Y., Fu, Z., Bustillo, J., Turner, J.A., Calhoun, V.D., Sui, J., 2020. Task-induced brain connectivity promotes the detection of individual differences in brain-behavior relationships. NeuroImage 207, 116370.

Kong, R., Li, J., Orban, C., Sabuncu, M.R., Liu, H., Schaefer, A., Sun, N., Zuo, X.N., Holmes, A.J., Eickhoff, S.B., Yeo, B.T.T., 2019. Spatial Topography of Individual-Specific Cortical Networks Predicts Human Cognition, Personality, and Emotion. Cereb Cortex 29, 2533–2551.

Kveraga, K., Ghuman, A.S., Bar, M., 2007. Top-down predictions in the cognitive brain. Brain Cogn 65, 145–168.

Li, J., Kong, R., Liegeois, R., Orban, C., Tan, Y., Sun, N., Holmes, A.J., Sabuncu, M.R., Ge, T., Yeo, B.T.T., 2019. Global signal regression strengthens association between resting-state functional connectivity and behavior. NeuroImage 196, 126–141.

Liégeois, R., Li, J., Kong, R., Orban, C., Van De Ville, D., Ge, T., Sabuncu, M.R., Yeo, B.T.T., 2019. Resting brain dynamics at different timescales capture distinct aspects of human behavior. Nature Communications 10, 2317.

Mansour, L.S., Tian, Y., Yeo, B.T.T., Cropley, V., Zalesky, A., 2021. High-resolution connectomic fingerprints: Mapping neural identity and behavior. NeuroImage 229, 117695.

Marek, S., Tervo-Clemmens, B., Calabro, F.J., Montez, D.F., Kay, B.P., Hatoum, A.S., Donohue, M.R., Foran, W., Miller, R.L., Feczko, E., Miranda-Dominguez, O., Graham, A.M., Earl, E.A., Perrone, A.J., Cordova, M., Doyle, O., Moore, L.A., Conan, G., Uriarte, J., Snider, K., Tam, A., Chen, J., Newbold, D.J., Zheng, A., Seider, N.A., Van, A.N., Laumann, T.O., Thompson, W.K., Greene, D.J., Petersen, S.E., Nichols, T.E., Yeo, B.T.T., Barch, D.M., Garavan, H., Luna, B., Fair, D.A., Dosenbach, N.U.F., 2020. Towards Reproducible Brain-Wide Association Studies. bioRxiv, 2020.2008.2021.257758.

Marewski, J.N., Schooler, L.J., 2011. Cognitive niches: an ecological model of strategy selection. Psychol Rev 118, 393–437.

Miller, K.L., Alfaro-Almagro, F., Bangerter, N.K., Thomas, D.L., Yacoub, E., Xu, J., Bartsch, A.J., Jbabdi, S., Sotiropoulos, S.N., Andersson, J.L., Griffanti, L., Douaud, G., Okell, T.W., Weale, P., Dragonu, I., Garratt, S., Hudson, S., Collins, R., Jenkinson, M., Matthews, P.M., Smith, S.M., 2016. Multimodal population brain imaging in the UK Biobank prospective epidemiological study. Nat Neurosci 19, 1523–1536.

Noble, S., Scheinost, D., Constable, R.T., 2019. A decade of test-retest reliability of functional connectivity: A systematic review and meta-analysis. NeuroImage 203, 116157.

Noble, S., Scheinost, D., Constable, R.T., 2021. A guide to the measurement and interpretation of fMRI test-retest reliability. Current Opinion in Behavioral Sciences 40, 27–32.

Noble, S., Spann, M.N., Tokoglu, F., Shen, X., Constable, R.T., Scheinost, D., 2017. Influences on the Test–Retest Reliability of Functional Connectivity MRI and its Relationship with Behavioral Utility. Cerebral Cortex 27, 5415–5429.

Nogueira, S., Sechidis, K., Brown, G., 2017. On the stability of feature selection algorithms. J. Mach. Learn. Res. 18, 6345–6398.

Pannunzi, M., Hindriks, R., Bettinardi, R.G., Wenger, E., Lisofsky, N., Martensson, J., Butler, O., Filevich, E., Becker, M., Lochstet, M., Kühn, S., Deco, G., 2017. Resting-state fMRI correlations: From link-wise unreliability to whole brain stability. NeuroImage 157, 250–262.

Park, S.A., Sestito, M., Boorman, E.D., Dreher, J.-C., 2019. Neural computations underlying strategic social decision-making in groups. Nature Communications 10, 5287.

Pervaiz, U., Vidaurre, D., Woolrich, M.W., Smith, S.M., 2020. Optimising network modelling methods for fMRI. NeuroImage 211, 116604.

Poldrack, R.A., Baker, C.I., Durnez, J., Gorgolewski, K.J., Matthews, P.M., Munafò, M.R., Nichols, T.E., Poline, J.B., Vul, E., Yarkoni, T., 2017. Scanning the horizon: towards transparent and reproducible neuroimaging research. Nat Rev Neurosci 18, 115–126.

Poldrack, R.A., Huckins, G., Varoquaux, G., 2020. Establishment of Best Practices for Evidence for Prediction: A Review. JAMA Psychiatry 77, 534–540.

Power, J.D., Barnes, K.A., Snyder, A.Z., Schlaggar, B.L., Petersen, S.E., 2012. Spurious but systematic correlations in functional connectivity MRI networks arise from subject motion. NeuroImage 59, 2142–2154.

Robinson, E.C., Jbabdi, S., Glasser, M.F., Andersson, J., Burgess, G.C., Harms, M.P., Smith, S.M., Van Essen, D.C., Jenkinson, M., 2014. MSM: a new flexible framework for Multimodal Surface Matching. NeuroImage 100, 414–426.

Rosenberg, M.D., Casey, B.J., Holmes, A.J., 2018a. Prediction complements explanation in understanding the developing brain. Nature Communications 9, 589.

Rosenberg, M.D., Finn, E.S., Scheinost, D., Papademetris, X., Shen, X., Constable, R.T., Chun, M.M., 2016. A neuromarker of sustained attention from whole-brain functional connectivity. Nat Neurosci 19, 165–171.

Rosenberg, M.D., Hsu, W.-T., Scheinost, D., Todd Constable, R., Chun, M.M., 2018b. Connectome-based Models Predict Separable Components of Attention in Novel Individuals. J Cogn Neurosci 30, 160–173.

Salimi-Khorshidi, G., Douaud, G., Beckmann, C.F., Glasser, M.F., Griffanti, L., Smith, S.M., 2014. Automatic denoising of functional MRI data: combining independent component analysis and hierarchical fusion of classifiers. NeuroImage 90, 449–468.

Schulz, M.-A., Yeo, B.T.T., Vogelstein, J.T., Mourao-Miranada, J., Kather, J.N., Kording, K., Richards, B., Bzdok, D., 2020. Different scaling of linear models and deep learning in UKBiobank brain images versus machine-learning datasets. Nature Communications 11, 4238.

Seguin, C., Tian, Y., Zalesky, A., 2020. Network communication models improve the behavioral and functional predictive utility of the human structural connectome. Netw Neurosci 4, 980–1006.

Shen, X., Finn, E.S., Scheinost, D., Rosenberg, M.D., Chun, M.M., Papademetris, X., Constable, R.T., 2017. Using connectome-based predictive modeling to predict individual behavior from brain connectivity. Nat Protoc 12, 506–518.

Shirer, W.R., Jiang, H., Price, C.M., Ng, B., Greicius, M.D., 2015. Optimization of rs-fMRI Pre-processing for Enhanced Signal-Noise Separation, Test-Retest Reliability, and Group Discrimination. NeuroImage 117, 67–79.

Smith, S.M., Beckmann, C.F., Andersson, J., Auerbach, E.J., Bijsterbosch, J., Douaud, G., Duff, E., Feinberg, D.A., Griffanti, L., Harms, M.P., Kelly, M., Laumann, T., Miller, K.L., Moeller, S., Petersen, S., Power, J., Salimi-Khorshidi, G., Snyder, A.Z., Vu, A.T., Woolrich, M.W., Xu, J., Yacoub, E., Ugurbil, K., Van Essen, D.C., Glasser, M.F., 2013. Resting-state fMRI in the Human Connectome Project. NeuroImage 80, 144–168.

Sui, J., Jiang, R., Bustillo, J., Calhoun, V., 2020. Neuroimaging-based Individualized Prediction of Cognition and Behavior for Mental Disorders and Health: Methods and Promises. Biol Psychiatry 88, 818–828.

Taxali, A., Angstadt, M., Rutherford, S., Sripada, C., 2021. Boost in Test–Retest Reliability in Resting State fMRI with Predictive Modeling. Cerebral Cortex 31, 2822–2833.

Tian, Y., Margulies, D.S., Breakspear, M., Zalesky, A., 2020. Topographic organization of the human subcortex unveiled with functional connectivity gradients. Nat Neurosci 23, 1421–1432.

Tibshirani, R., 1996. Regression Shrinkage and Selection via the Lasso. Journal of the Royal Statistical Society. Series B (Methodological) 58, 267–288.

Tibshirani, R.J., 2013. The lasso problem and uniqueness. Electronic Journal of Statistics 7, 1456-1490, 1435.

Van Essen, D.C., Smith, S.M., Barch, D.M., Behrens, T.E.J., Yacoub, E., Ugurbil, K., 2013. The WU-Minn Human Connectome Project: An overview. NeuroImage 80, 62–79.

Varoquaux, G., 2018. Cross-validation failure: Small sample sizes lead to large error bars. NeuroImage 180, 68–77.

Wang, D., Li, M., Wang, M., Schoeppe, F., Ren, J., Chen, H., Öngür, D., Brady, R.O., Baker, J.T., Liu, H., 2020. Individual-specific functional connectivity markers track dimensional and categorical features of psychotic illness. Mol Psychiatry 25, 2119–2129.

Weintraub, S., Dikmen, S.S., Heaton, R.K., Tulsky, D.S., Zelazo, P.D., Bauer, P.J., Carlozzi, N.E., Slotkin, J., Blitz, D., Wallner-Allen, K., Fox, N.A., Beaumont, J.L., Mungas, D., Nowinski, C.J., Richler, J., Deocampo, J.A., Anderson, J.E., Manly, J.J., Borosh, B., Havlik, R., Conway, K., Edwards, E., Freund, L., King, J.W., Moy, C., Witt, E., Gershon, R.C., 2013. Cognition assessment using the NIH Toolbox. Neurology 80, S54–64.

Woolrich, M.W., Jbabdi, S., Patenaude, B., Chappell, M., Makni, S., Behrens, T., Beckmann, C., Jenkinson, M., Smith, S.M., 2009. Bayesian analysis of neuroimaging data in FSL. NeuroImage 45, S173–186.

Yarkoni, T., Westfall, J., 2017. Choosing Prediction Over Explanation in Psychology: Lessons From Machine Learning. Perspect Psychol Sci 12, 1100–1122.

Yeo, B.T., Krienen, F.M., Sepulcre, J., Sabuncu, M.R., Lashkari, D., Hollinshead, M., Roffman, J.L., Smoller, J.W., Zollei, L., Polimeni, J.R., Fischl, B., Liu, H., Buckner, R.L., 2011. The organization of the human cerebral cortex estimated by intrinsic functional connectivity. J Neurophysiol 106, 1125–1165.

Zalesky, A., Fornito, A., Bullmore, E.T., 2010. Network-based statistic: identifying differences in brain networks. NeuroImage 53, 1197–1207.

## Reference

Hoerl AE, Kennard RW. 1970. Ridge Regression: Biased Estimation for Nonorthogonal Problems. Technometrics 12:55–67.

Li J, Kong R, Liegeois R, Orban C, Tan Y, Sun N, Holmes AJ, Sabuncu MR, Ge T, Yeo BTT. 2019. Global signal regression strengthens association between resting-state functional connectivity and behavior. Neuroimage 196:126–141.

Nocedal J, Wright SJ. 2006. Numerical Optimization. New York: Springer.

Shen X, Finn ES, Scheinost D, Rosenberg MD, Chun MM, Papademetris X, Constable RT. 2017. Using connectome-based predictive modeling to predict individual behavior from brain connectivity. Nat Protoc 12:506–518.

Tibshirani R. 1996. Regression Shrinkage and Selection via the Lasso. Journal of the Royal Statistical Society Series B (Methodological) 58:267–288.

Wright SJ, Nowak RD, Figueiredo MAT. 2009. Sparse Reconstruction by Separable Approximation. IEEE Transactions on Signal Processing 57:2479–2493.

